# Cnr2 is important for Ribbon Synapse Maturation and Function in Hair Cells and Photoreceptors

**DOI:** 10.1101/2020.08.18.253120

**Authors:** Luis Colon-Cruz, Roberto Rodriguez-Morales, Alexis Santana-Cruz, Juan Cantres-Velez, Aranza Torrado-Tapias, Guillermo Yudowski, Robert Kensler, Bruno Marie, Shawn Burgess, Olivier Renaud, Gaurav K. Varshney, Martine Behra

## Abstract

The role of the cannabinoid receptor 2 (CNR2) is still poorly described in sensory epithelia. We found strong *cnr2* expression in hair cells (HCs) of the inner ear and the lateral line (LL), a superficial sensory structure in fish. Next, we demonstrated that sensory synapses in HCs were severely perturbed in zebrafish larvae lacking cnr2. Appearance and distribution of presynaptic ribbons and calcium channels (Ca_v_1.3) were profoundly altered in mutant animals. Clustering of membrane-associated guanylate kinase (MAGUK) in post-synaptic densities (PSDs) was also heavily affected, suggesting a role for cnr2 for maintaining the structure of the sensory synapse. Furthermore, vesicular trafficking in HCs was strongly perturbed suggesting a retrograde action of the endocannabinoid system (ECs) mediated via cnr2 that was modulating HC mechanotransduction. We found similar perturbations in retinal ribbon synapses. Finally, we showed that larval swimming behaviors after sound and light stimulations were significantly different in mutant animals. Thus, we propose that cnr2 is critical for the processing of sensory information in the developing larva.

## INTRODUCTION

The endocannabinoid system (ECs) is a well conserved modulator of almost all physiological systems, including the central and peripheral nervous systems [1]. Endocannabinoids mostly serve as retrograde messenger at various types of synapses throughout the brain, where they are synthesized in response to post-synaptic activation and bind pre-synaptic cannabinoid receptor 1 (CNR1) which in turn inhibits classical neurotransmitter release [2]. CNR1 expression and function was extensively described in the brain [3] but also in several peripheral organs [4]. By contrast, expression, and function of the second well characterized cannabinoid receptor (CNR2) remained more elusive until recently, in part due to interspecies differences at the genomic and tissue/organ expression levels [5]. The previously accepted notion that CNR2 was mostly restricted to the immune system has been repeatedly challenged [6], and expression in the mammalian [7] and fish brain [8] as well as associated behavioral changes were demonstrated. In sensory organs, *Cnr2* expression was described in several cell types like photoreceptors of the adult retina in various mammals [9] as well as in fish [10]. More recently, Cnr2 expression was found in several cell types of the rodent inner ear like sensory hair cells (HCs) of the Organ of Corti [11, 12]. However, little is known about the developmental expression pattern or the role of cnr2 in those epithelia.

HCs are sensory mechanoreceptors which are found in the auditory part of the mammalian inner ear in the floor of the cochlear canal (= Organ of Corti), but also in sensory patches of the vestibular part, which is common to all vertebrates, and is comprised of 3 cristae and 2 maculae, that insure static and dynamic equilibrium [13]. In fish, in addition to a well conserved inner ear, there is an evolutionary linked superficial sensory organ named the lateral line (LL) that is composed of sensory patches called neuromasts (NMs) [14]. NMs are stereotypically and symmetrically distributed on each side of the animal’s head (= anterior LL, aLL), trunk and tail (= posterior LL, pLL). In zebrafish, both the ear and the LL appear early in the embryo and mature rapidly, becoming functional in the 3 to 5day post-fertilization (dpf) larva in a rostral to caudal gradient for the pLL [15]. In mature HCs, the apical bundle of cilia are deflected by sound waves and head movements in the inner ear, and by water movements in the LL. The opening of mechanical ion channels triggers a graded membrane depolarization [16], that in turn activates the voltage gated calcium channels in the latero-basal synapses, thus creating a calcium influx, that will drive the fusion od synaptic vesicles and release of neurotransmitter (glutamate) onto the innervating afferent fibers [17]. HCs must transmit sound over a dynamic range of several orders of magnitude similar to how photoreceptors transmit light signals, meaning that changes of intensity in the stimulus are encoded by adjusting the tonic (sustained) rate of neurotransmitter release [18]. A phasic (transient) rate of vesicle release was also demonstrated in retinal cones exposed to light when detecting variation in light intensity [19, 20]. Graded synaptic output requires the release of up to several thousand synaptic vesicles/second. This is made possible by signature structures that are exclusively found in sensory receptor cell synapses that are called ribbons or dense bodies.

Ribbons are anchored to the basolateral membrane directly adjacent to synaptic cluster of L-type voltage gated calcium channels and are surrounded by tethered glutamate-filled vesicles [21, 22]. The ribbon itself is a dynamic electron dense structure that can adopt different shapes, typically forming plates in retinal cells [18], and spheroids in HCs [22, 23]. The main constituent of ribbons is the RIBEYE protein that is encoded by the *CTBP2* gene with a unique N-terminal A domain that does not share homology with any other known protein, and a C-terminal B domain identical to the CTBP2 protein [24]. Furthermore, both the size and number of ribbons per sensory cell vary depending on the species [25], the cell type [25], the position in the sensory epithelium [26], the developmental stage [27, 28], and the sensory activity [29]. In the LL of zebrafish larva, mature HCs arise as early as 3dpf, in which three to four spherical ribbons (diameter ^~^200-300nm) neatly arranged in the basolateral portion can be visualized [24, 30–32]. Tethered spherical (diameter ^~^20-30nm) glutamate-filled vesicles surround the ribbon body. A subset in direct contact with the cytoplasmic membrane is docked and represents the ready to release pool (RRP) [22, 25, 33]. Originally thought to work as a conveyor belt the ribbon is now more viewed as a safety belt slowing down and organizing compound fusion of vesicles at the synapses [21, 34].

A number of additional proteins have been described at the ribbon synapse, some of them are common to classical synapses, and others exclusive (for review[27] [35]) like the highly-conserved L-type calcium channels (Ca_v_1.4 in the retina and Ca_v_1.3 in HCs). Absence of Ca_v_1.3 causes profound deafness in humans [36], mice [37], and zebrafish [38]. Furthermore, a close relationship between Ca_v_1.3 distribution and ribbon size and position was carefully characterized [31, 39, 40] confirming the crucial role of both for proper synaptic exo- and endocytosis [40, 41]. A classical approach to study synaptic vesicular trafficking is the application of FM 1-43, a styryl probe that fluoresces brightly when taken up in a membrane, that was widely used in a variety of preparations like neuro-muscular junctions [42–45], cultured hippocampal neurons [46] and retinal bipolar cells [47]. However, in HCs, FM 1-43 rapidly penetrates the apical ciliated region via two proposed mechanisms: apical cuticular endocytosis [48–52], and passive diffusing through mechanotransduction channels (MET) [53–56] and ion channels [57]. Thus, FM 1-43 will also report constitutive membrane trafficking [58] which can potentially mask synaptic vesicle trafficking [59, 60]. Nevertheless, FM 1-43 was extensively used for assessing mechano-transduction during development [38, 48, 61, 62], intracellular apico-basal trafficking [63] [64] and synaptic recycling in HCs [65].

We found that the *cnr2* gene was expressed in all HCs of the inner ear and the LL, starting at 3day post-fertilization (dpf) onward. In *cnr2* loss of function animals (*cnr2^upr1^*) that we had previously generated [8], notably HCs in NMs of the LL were not strongly affected in their development, regeneration, or innervation. However, we found profound perturbation in the size and distribution of synaptic ribbons which was more evident in mature HCs. Furthermore, we showed alteration in the distribution of Ca_v_1.3 channels, and we demonstrated that the alignment of pre- and post-synaptic elements in afferent dendrites was compromised. Using TEM and live FM 1-43 imaging, we showed that the trafficking of neurotransmitter vesicles was affected, suggesting a putative cnr2 mediated retrograde activity of the ECs in HCs. We further observed perturbation of the ribbon synapses in the retina of *cnr2^upr/upr1^* animals, raising the possibility of a mechanism that is common to all sensory synapses. Finally, we established that mutant larvae were more sensitive to sound stimulation when also exposed to light, and conversely more sensitive to light when previously exposed to sound, thus providing a link between sensory tasks in a cnr2-deficient context and illustrating the physiological relevance of our findings.

## RESULTS

### Cnr2 is strongly expressed in hair cells (HCs) of the LL and the inner ear

To define the *cnr2* expression pattern during development, we performed whole mount *in situ* hybridization (WISH) with an antisense probe against *cnr2* in 3 and 5day post-fertilization (dpf, n= 30/stage) wild type animals. At 3dpf (Figure 1) and 5dpf (not shown), we found a strong hair cell (HC)-specific expression in all neuromasts (NMs) of the lateral line (LL, Figure 1A, B, D, E, F, G and H), as well as in all sensory patches of the inner ear, namely the maculae (1C, #) and cristae (1C, *).

**Figure 1.**
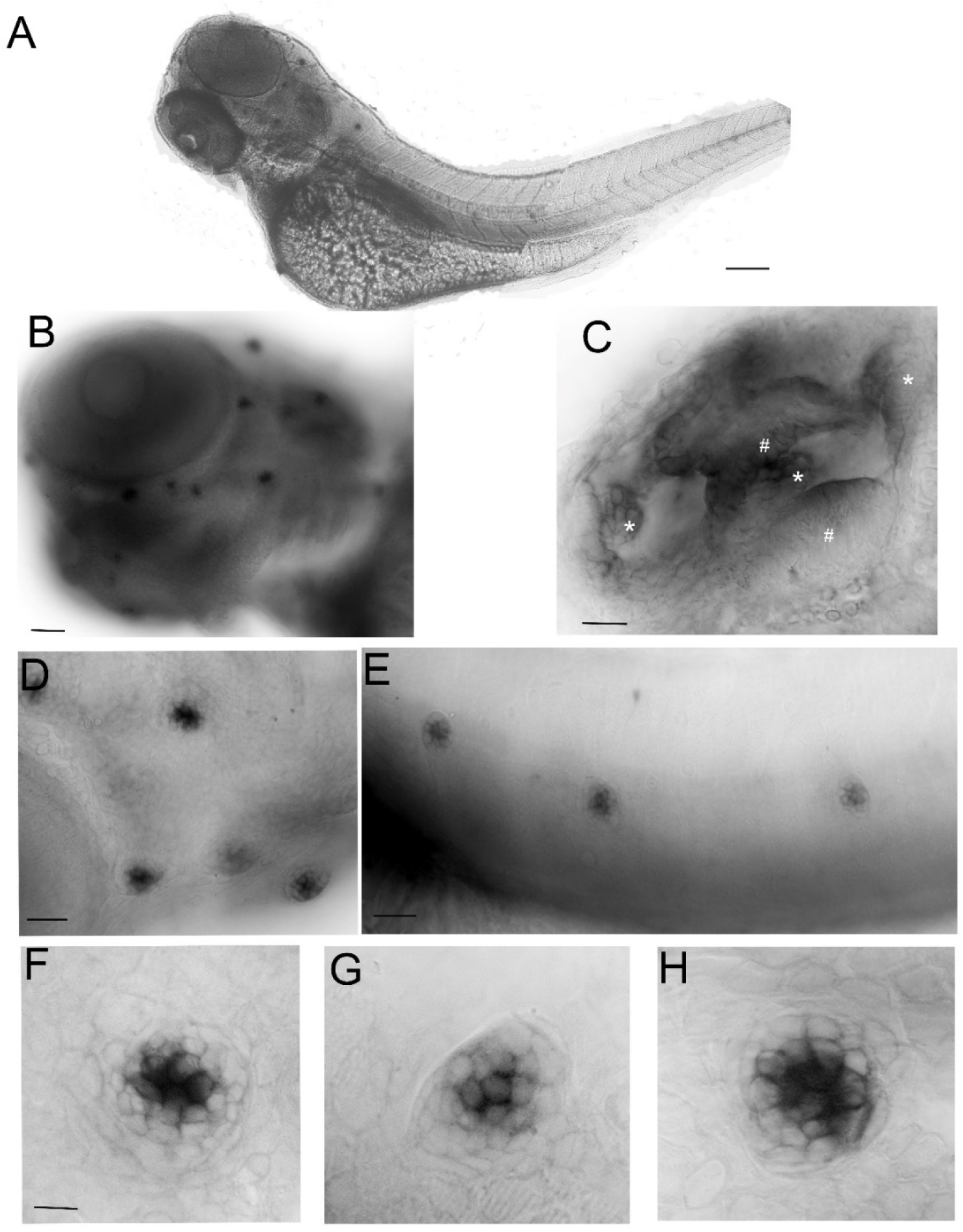
Whole mount in situ hybridization (WISH) with an antisense probe against *cnr2* in 3day post-fertilization (dpf) wild type larvae. (**A**). Lateral view of a whole larva. (**B**). Magnification of the head region showing head neuromasts (NMs) and inner ear staining. (**C**). Inner ear showing labelling in all three cristae (*) and two maculae (#). (**D**). Magnification of cranial NMs in the anterior lateral line (LL) (**E**). Magnification of trunk NMs in the posterior LL. (**F, G**, and **H**). Individuals NMs show *cnr2* expression that is restricted to the centrally located HCs. Scale bars: in A = 1500 microns; in B = 300 microns; in C = 50 microns; in D = 20 microns; in E = 150 microns; in F, G and H = 25 microns.

To elucidate the putative role of cnr2 in those sensory epithelia, we used the stable loss-of-function mutant line (*cnr2^upr1^*) that we had previously generated [8], to monitor HC development and regeneration in the LL in the absence of cnr2 (Figure1-supplement 1). We visualized functional HCs using FM 1-43 [48]. We counted HCs in successive developmental stages (1-sup 1A and 1B), namely 2, 3, 4, 5, 6, and 7 dpf (with a minimum of n=22/genotype/stage) wild type and *cnr2^upr1/upr1^* larvae. The overall development of the LL and the gross morphology of NMs seemed unaffected (1-sup 1A), but the average HCs number/NM was slightly but significantly reduced in mutant larvae starting at 3dpf and onward (1-sup 1B, light grey bars). Next, we triggered synchronized HC-destruction using copper treatments in 5dpf wild type and mutant larvae and counted regenerating HCs at 0, 24, 48 and 72hour post-treatment (hpt, 1-sup 1C, n=23/genotype/hpt). We found that HC regeneration was not significantly affected until 48hpt and slightly decreased thereafter. Taken together, *cnr2* was strongly expressed in HCs of the LL but when absent, both HC development and regeneration were only moderately affected, suggesting cnr2 was not indispensable for either.

### Cnr2 expression affects ribbon synapses in mature HCs of the LL and the inner ear

Next, we focused on the LL and assessed efferent and afferent innervation of NMs. We immunolabelled 5dpf wild type and *cnr2^upr1/upr1^* larvae with two classical antibodies (Abs) (1) Znp1 which stains motor neurons projections and terminals, and (2) Myosin7 which labels HCs (Figure 1-Supplement 2, n=15/genotype). We found no overt differences in the efferent innervation of NMs (1-sup2A). In parallel, we immunolabelled sensory neurons projections using Zn12 another classical Ab, and co-labelled ribbons synapses in HCs with an anti-Ribeye b Ab (1-sup2B n= 12/genotype). The overall sensory innervation appears intact (Zn12 in magenta) however, the Ribeye b staining (green) was strongly perturbed in mutant NMs, suggesting that ribbon synapses were defective in the absence of proper cnr2 expression.

To further analyze the sensory synapses, we co-immunolabelled 5dpf wild type (n= 20) and mutant larvae (n=23) with Abs against Ribeye b, and against the scaffolding protein membrane-associated guanylate kinase (Maguk) which forms foci in post-synaptic densities (PSDs) of sensory neurites [30, 40] (Figure 2). In wild type NMs (first and second rows in A and B) in both the anterior LL (aLL, Figure.2A) and the posterior LL (pLL, 2B), we found that Ribeye b puncta (green) appeared neatly organized, and localized to the laterobasal portion of HCs (more visible in 90^0^ rotation of the acquired images, second rows in A and B). On average, we found 3 to 4 evenly shaped and sized puncta/HC as previously described [31, 66]. By contrast, in NMs of cnr2-KOs (third and fourth rows in A and B), we found much less Ribeye b puncta, which were highly variable in shape and size, unorganized and not always localized to the basal portion of HCs. Furthermore, in the PSDs of wild type NMs and as previously described, we found focalized Maguk staining (magenta in first and second rows in 2A and 2B) which was mostly located in close vicinity to the Ribeye b puncta, thus forming bi-labelled clusters (inserts on extreme right). By contrast, in mutant NMs, Maguk staining was often weak and diffuse with few foci which when present were rarely in the close vicinity of Ribeye b puncta. Taken together, this was strongly suggesting that the sensory synapses were perturbed both in the presynaptic zone in HCs but also in the post-synaptic dendrites, and that the alignment of synaptic elements was challenged.

**Figure 2.**
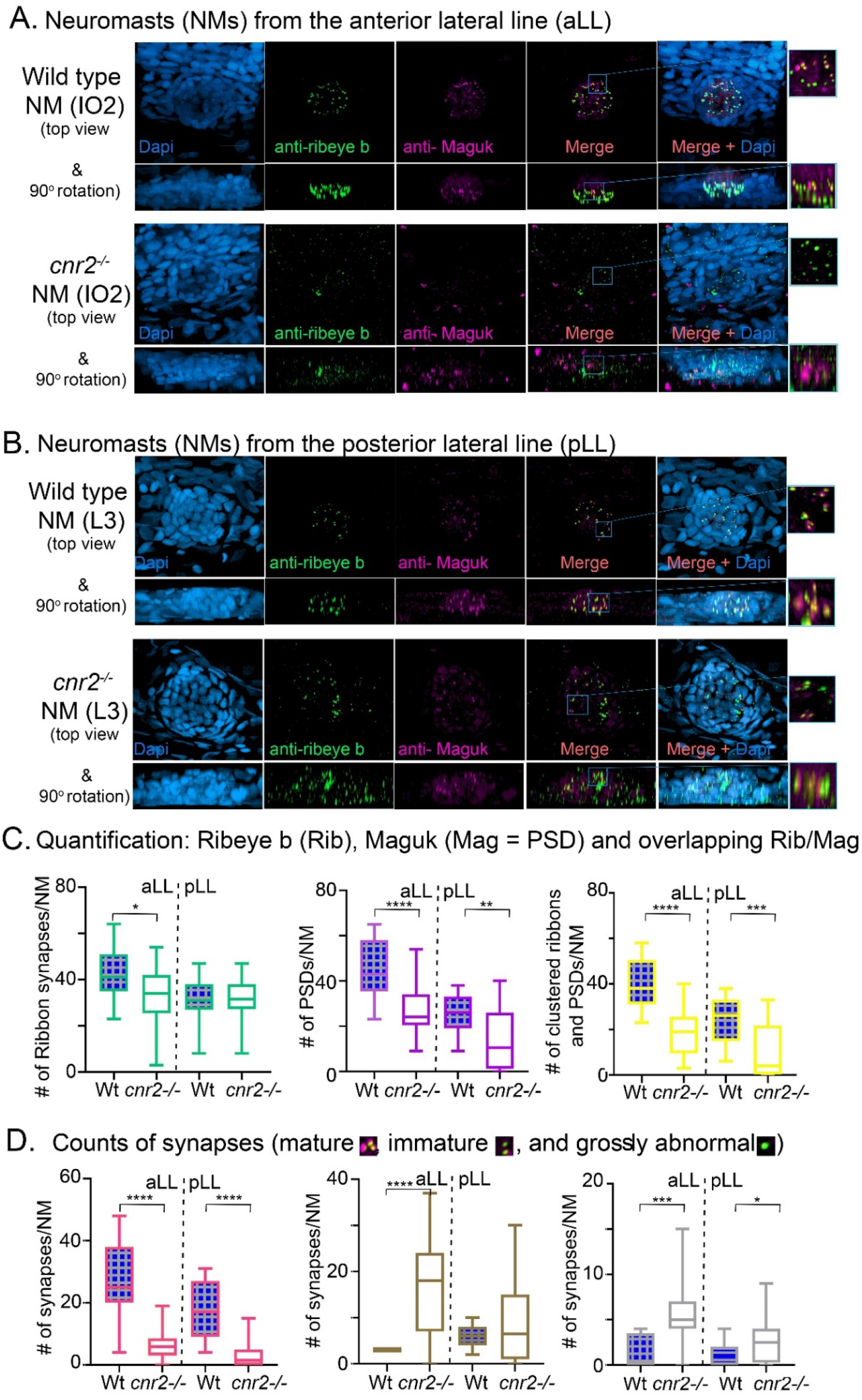
Immunofluorescent labelling of the ribbon synapses (Ribeye b) in HCs and the post-synaptic densities (PSDs, Maguk) in afferent dendrites in neuromasts (NMs) of wild type (top panels) and *cnr2^upr1/upr1^* (bottom panels) 5dpf larvae. (**A**). Cranial NMs (IO2) from the anterior lateral line (aLL) in top views (top and middle lanes) and in 90° image rotations. Each respective region (blue squares) are magnified in the corresponding inserts (right panels). (**B**). Trunk NMs (L3) from the posterior lateral line (pLL) presented as in A. (**C**). Individual staining (= puncta) quantification of Ribeye b (left panel), Maguk (central panel) and the overlapping staining as illustrated in the inserts above (right panel). The aLL and pLL NMs were counted separately. (**D**). Visual assessment of the maturity of synapses was performed based on the size, shape and proportional overlap of Ribeye b/Maguk (Rib/Mag) staining and counted as mature (left panel), immature (central panel) or grossly abnormal (right panel). Scale bars: in A = 20 microns in left panel and = 1 micron in insert. In C and D: whisker boxes show all values with minimum and maximum. Significance is represented as * (* p < 0.05; ** p < 0.001; *** p < 0.003; and **** p < 0.0001)

Next, we quantified Ribeye b puncta, Maguk foci and the overlapping of both in wild type (n=4) and *cnr2^upr1/upr1^* larvae (n= 7) in NMs of the aLL (NM_WT_ n = 17; NM_cnr2_ n= 17) and pLL (NM_WT_ n = 19; NM_cnr2_ n= 20). For Ribeye b, we counted all puncta regardless of size (2C, left graph) and found that the number of ribbon synapses (RS) was significantly decreased in mutant NMs in the aLL (RS/NM_wt_ = 43.35 vs. RS/NM_cnr2_ = 33.94, p = 0.0256), but not in the pLL (RS/NM_wt_ = 30.99 vs. RS/NM_cnr2_ = 31.30, p = 0.8964), possibly because trunk and tail NMs were less mature than cranial NMs. For Maguk, we counted all foci regardless of the size (2C, middle graph), and found that the number of PSDs was significantly reduced in all NMs of aLL (PSD_WT_/NM = 45.47 vs. PSDcnr2/NM = 26.59, p < 0.0001), and pLL (PSD_WT_/NM = 25.74 vs. PSD_cnr2_/NM = 14.00, p = 0.0037). Finally, we scored all overlapping staining of Ribeye b and Maguk regardless of the respective size of either (2C, right graph) and we found that the number of RS clustered with PSDs was greatly reduced in aLL (RS/PSD_WT_/NM = 40.24 vs. RS/PSD_cnr2_/NM = 18.82, p < 0.0001) and in pLL (RS/PSD_WT_/NM = 23.95 vs. RS/PSD_cnr2_/NM = 10.40, p < 0.0003) of cnr2-KOs animals. Furthermore, only in wild type NMs did we find the previously described mature aligned synapses which are forming evenly-sized, balanced clusters [30, 31]. In the mutant NMs, when we did find bi-labelled clusters they appeared mostly uneven in size and labelling, presumably reflecting mostly immature synapses (inserts in right columns). Thus, we next subdivided the co-labelled clusters into three categories, (1) mature (2D, left graph), (2) immature when either staining in the cluster was strongly unbalanced (2D, middle graph), (3) and grossly abnormal when either staining was obviously dysmorphic (2D, right graph). Mature and normal appearing synapses (S) were clearly less abundant in mutant NMs of the aLL (S_WT_/NM = 26.65 vs. S_cnr2_/NM = 6.50, p < 0.0001) and pLL (S_WT_/NM = 17.95 vs. S_cnr2_/NM = 3.80, p < 0.0001). Immature synapses (IS) were rarely found in wild type NMs from the aLL, but their number was strongly increased in mutant animals (IS_WT_/NM = 3.00 vs. IS_cnr2_/NM = 16.24, p < 0.0001). Notably, in the pLL, IS numbers were not significantly different (IS_WT_/NM = 6.316 vs. IS_cnr2_/NM = 8.800, p < 0.2472), possibly reflecting the ongoing antero-posterior maturation of NMs in the LL. Finally, we only found abnormal synapses (AS) in aLL (AS_WT_/NM = 1.41 vs. AS_cnr2_/NM = 5.41, p = 0.0005) and pLL (AS_WT_/NM = 1.16 vs. AS_cnr2_/NM = 2.90, p = 0.0192) mutant NMs. Taken together, in the absence of cnr2 expression all HCs of the LL presented strongly altered Ribeye b staining as well as perturbed Maguk staining in the PSDs in afferent neurites. Alterations were generally stronger in the more mature NMs of the aLL, suggesting that cnr2 is possible involved in the maturation of ribbon synapses in HCs.

To verify if HCs of sensory epithelia in the inner ear were also affected, we imaged and analyzed Ribeye b and Maguk staining in maculae (Figure 3) and cristae (not shown) of wild type (two top rows) and cnr2-KOs larvae (two bottom rows). Maculae at this stage, followed closely the curved shape of the inner ear displaying two opposing walls (see supplemental movie in Figure3-Supplement1) with most mature synapses on one of the two walls, as evidenced by more Ribeye b and Maguk overlapping clusters (see supplemental movie in Figure3-Supplement1-Wt). This apparent maturation gradient was also found in mutant maculae, but overall, both Ribeye b and Maguk staining appeared irregular. Notably, Ribeye b puncta (green) were more uneven in size while Maguk foci (magenta) were smaller and fewer, and the overlapping clusters were strongly reduced in number and size in cnr2-KO larvae (see magnified views in inserts on the right). Therefore, absence of cnr2 seemed also required for proper maturation of ribbon synapses in HCs of the inner ear, suggesting that the cnr2-dependent regulatory mechanism was common to all HCs in zebrafish larva.

**Figure 3.**
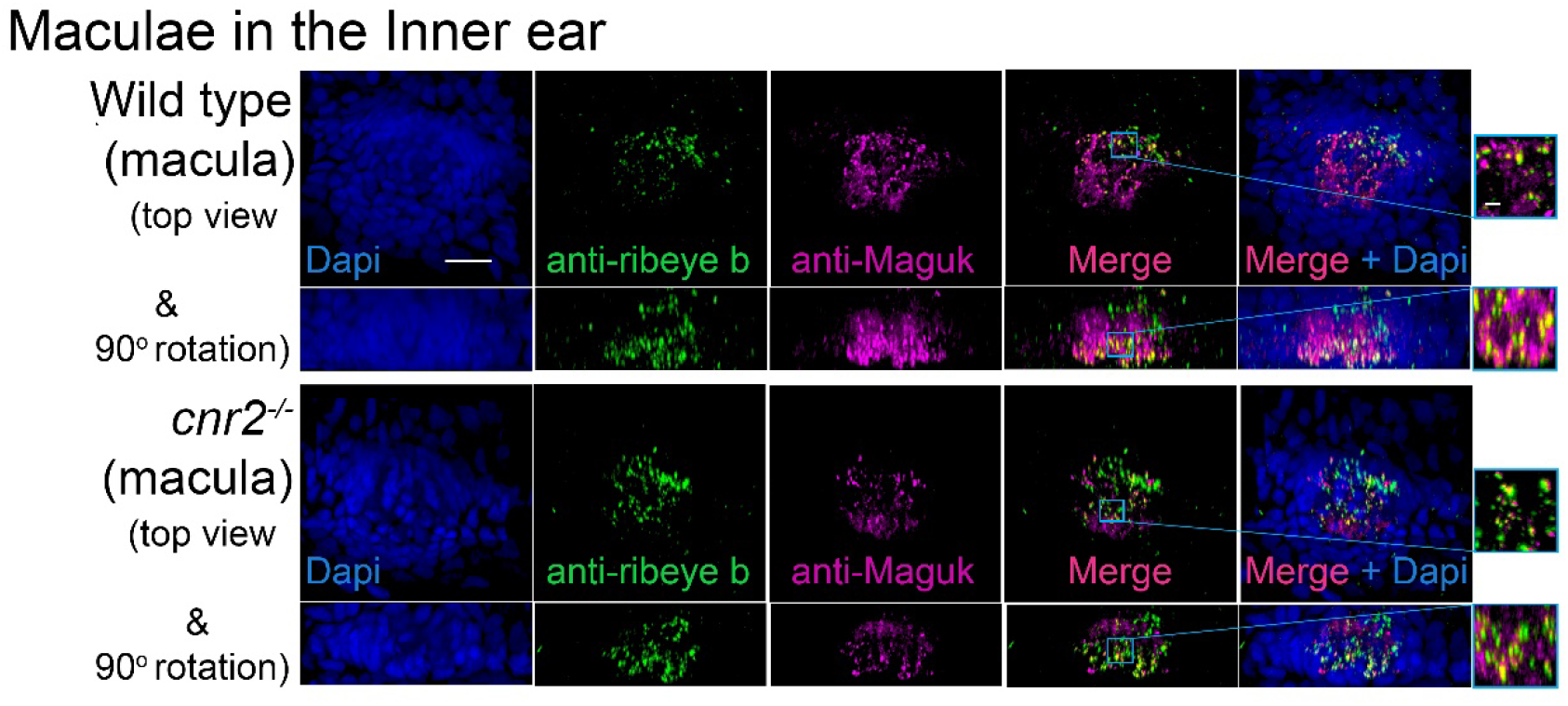
Immunofluorescence labelling of ribbon synapses (Ribeye b) and post synaptic densities (PSDs, Maguk) in 5dpf wild type (top panels) and *cnr2^upr1/upr1^* (bottom panels) inner ear maculae. Macula are stained with anti - Ribeye b (green) and anti-Maguk (magenta) Abs, in top views (top and middle rows) and in 90° image rotations (second and last rows). Magnification of the depicted region (blue squares) are presented in the corresponding inserts (right panels). Scale bars: in A= 20 microns in left panel and 1 micron in insert. **Supplemental movies:** Figure3-Supplement1-Rib-Maguk-macula Wt and Figure 3-Supplement1-Rib-Maguk-macula-cnr2

### Cnr2 controls presynaptic calcium channels (Ca_v_1.3) distribution in all HCs

To explore if Ca_v_1.3 distribution was affected in HCs of mutant larvae, we co-immunolabelled 5dpf wild type (n= 20) and mutant larvae (n=23) with Abs against Ca_v_1.3, and Maguk. We imaged the inner ear epithelia, namely cristae (Figure 4) and maculae (not shown). As described above, we found that Maguk staining was much weaker and more diffuse in mutant cristae (magenta in two lower panels) when compared to wild type cristae (two top panels). Strikingly, Ca_v_1.3 staining (green) was also strongly diminished, and there was no alignment between pre- and post-synaptic immunofluorescence in contrast to what we observed in wild type cristae (compare magnified regions (blue squares) in inserts in right columns). Thus, the Ca_v_1.3 distribution was profoundly altered in mutant inner ear epithelia.

**Figure 4.**
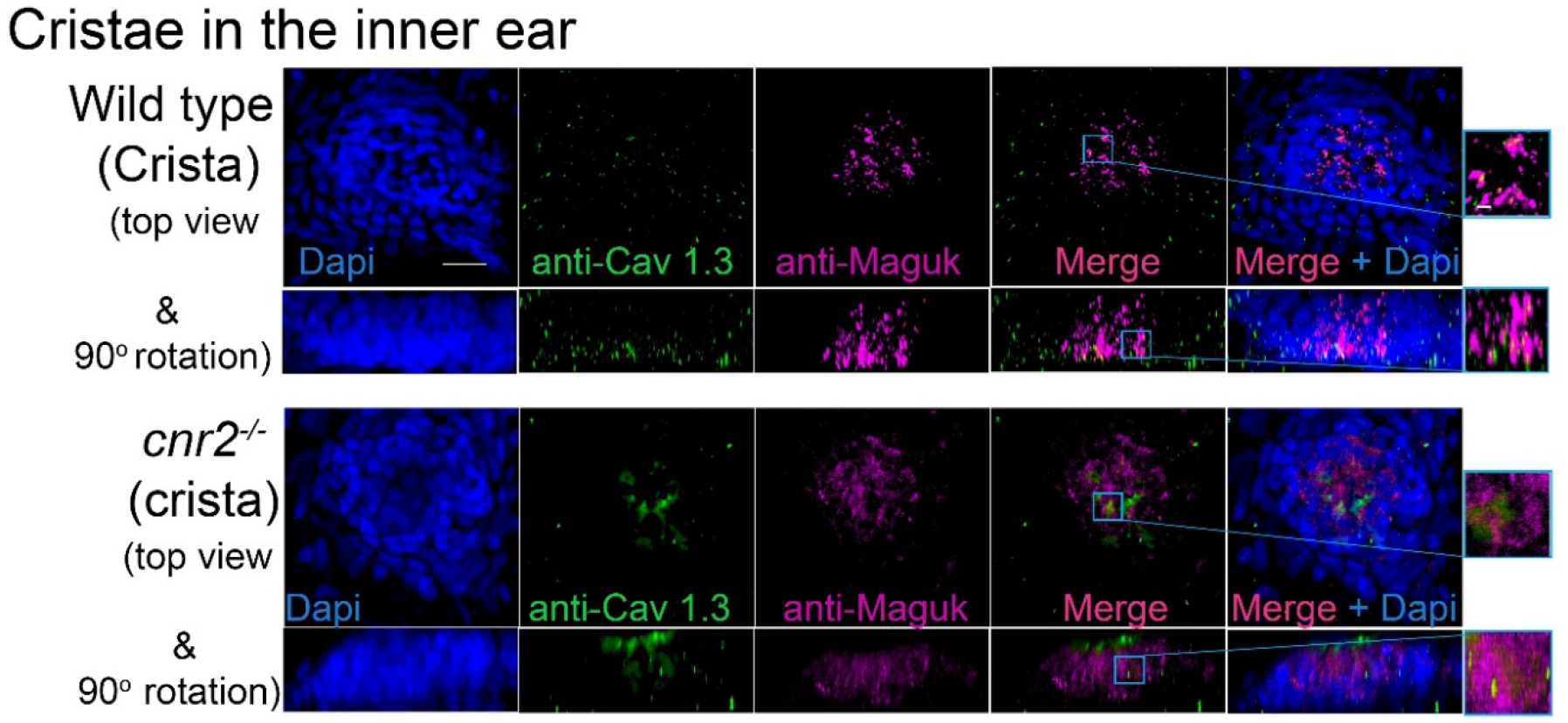
Immunofluorescence labelling of pre-synaptic calcium channels (Ca_v_1.3) and post-synaptic densities (PSDs, Maguk) countered stained with Dapi in HCs of 5dpf wild type (top panels) and *cnr2^upr1/upr1^* (bottom panels) inner ear cristae. Top views (top and middle rows) and 90° image rotations (second and last rows) of cristae that were stained with anti-Ca_v_1.3 (green) and anti-Maguk (magenta). Magnification of the depicted region (blue squares) are shown in the corresponding inserts (right panels). Scale bars: in = 20 microns in left panel and 1 micron in insert.

Next, we examined NMs of the aLL (not shown) and the pLL (Figure 5) and found that Maguk (magenta), and Ca_v_1.3 (green) staining were strongly altered. Taken together, the distribution of the Ca_v_1.3 channels was strongly perturbed in HCs of the inner ear epithelia as well as the LL in the absence of cnr2 expression.

**Figure 5.**
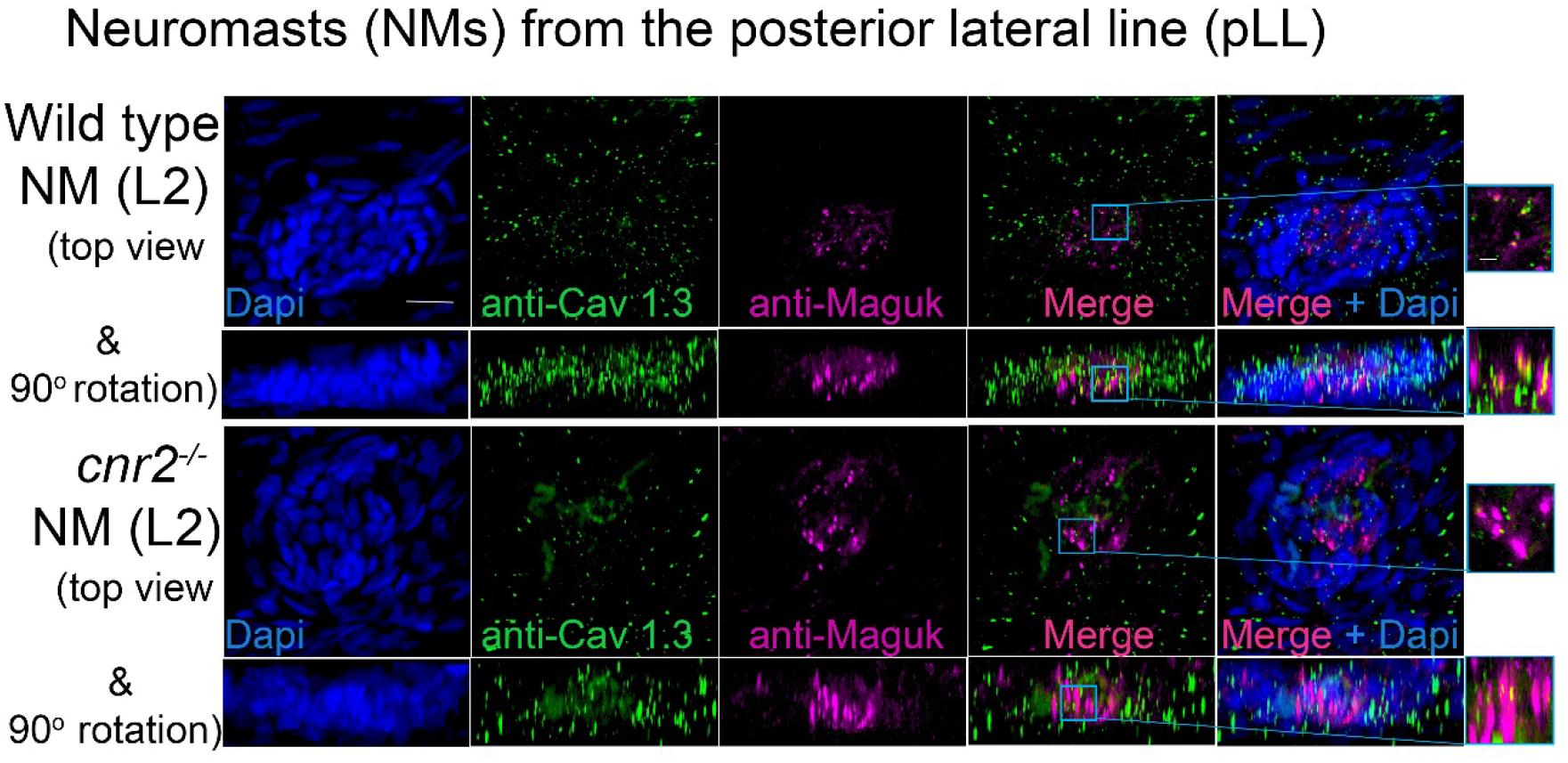
Immunofluorescent labelling of pre-synaptic calcium channels (Ca_v_1.3) and post-synaptic Maguk in PSDs counterstained with Dapi in wild type (two top panels) and *cnr2^upr1/upr1^* (two bottom panels) of 5dpf larvae neuromasts (NMs). Top views (first and third lanes) and 90° image rotations (second and fourth rows) of trunk NMs (L2) immunolabelled with Abs against Ca_v_1.3 (green) and Maguk (magenta). Each respective region (blue squares) are magnified in the corresponding inserts (right panels). Scale bars: 20 microns in left panel and 1 micron in insert.

### Cnr2 modulates distribution of neurotransmitter (glutamate) vesicles in HCs

To examine the ultra-structure of ribbons synapses and to verify if the distribution and appearance of neurotransmitter vesicle was also affected, we analyzed NMs of 5dpf wild type (n=3) and cnr2 homozygote (n=3) larvae using TEM (Figure 6). In all HCs of wild type (6A, C, E and H) and mutant (6B, D, F, G, I and J) NMs, we located ribbon synapses in the basolateral walls of the HCs and in close vicinity to innervating dendrites (red stars). At higher magnification and as expected, in wild type HCs we repeatedly found ribbon synapses which were regular in shape and size (two representative examples are shown in 6E and H). Likewise, the ribbon bodies were surrounded by regularly shaped and sized (^~^25nm) spherical vesicles (6E, white arrowheads). All were tethered at a constant distance (^~^10nm) and docked vesicles were also distinguishable (6H, white arrows). By contrast, in the mutant HCs, all the ribbon synapses that we found were presenting electron dense cores that were bigger and misshapen (representative illustrations are shown in 6F, G, I and J). Strikingly, the surrounding vesicles were often not spherical, highly variable in size and shape (magenta arrowheads) and at variable distance from the dense ribbon core. However, in most mutant HCs, the PSD was still clearly visible (blue arrows) and so were docked vesicles (magenta arrows in F, I and J), suggesting at least partial synaptic function. Taken together, the ultra-structure of the ribbon synapses and the surrounding vesicles was profoundly perturbed in animals lacking cnr2, pointing to an important role in the ultra-structural organization at the sensory synapse.

**Figure 6.**
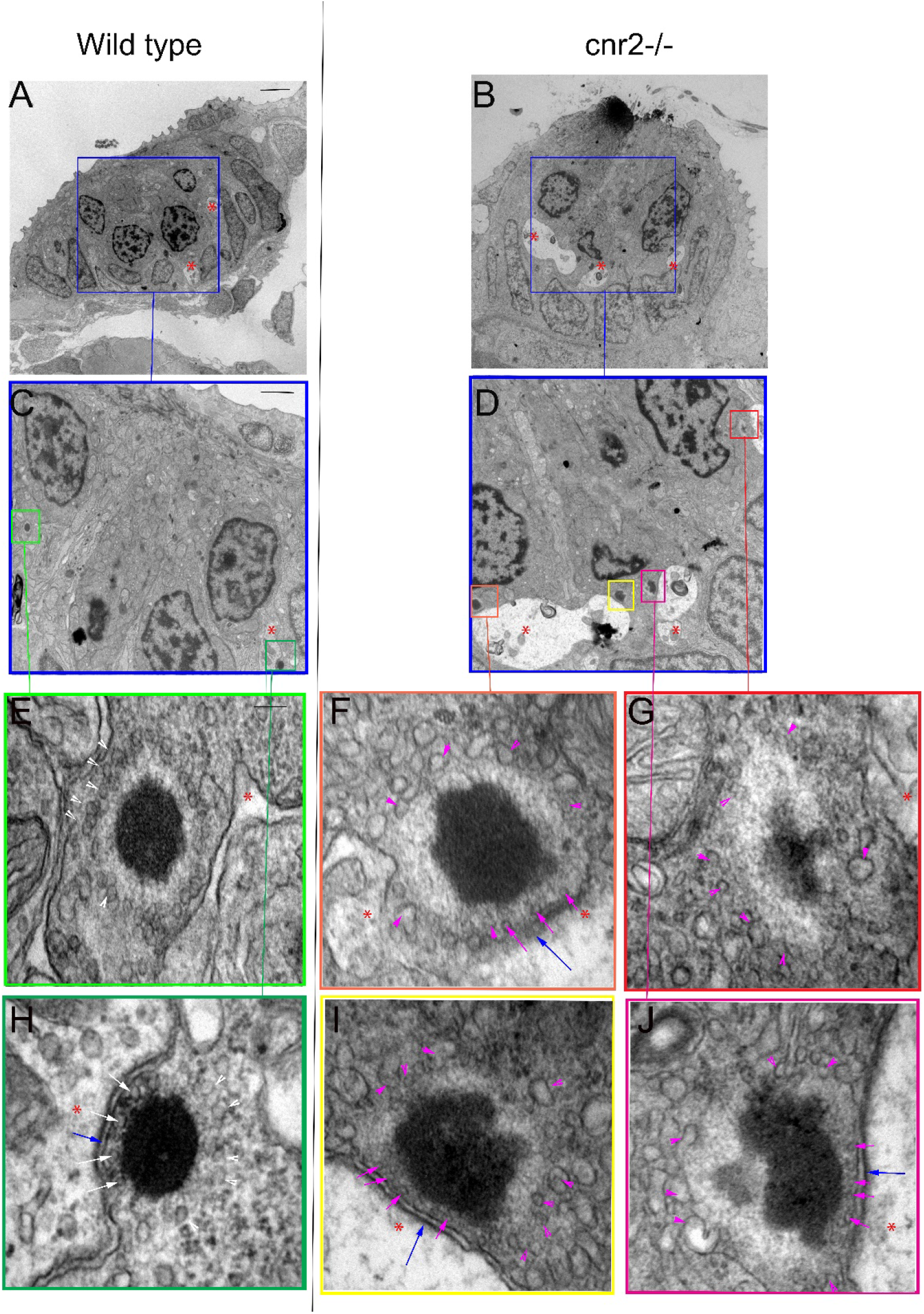
Transmission Electron microscopy (TEM) images in neuromasts (NMs) of 5dpf wild type and *cnr2^upr1/upr1^* mutant larvae. (**A** and **B**). Lower magnification showing the entire section through a wild type and a *cnr2^upr1/upr1^* NM respectively, with central ciliated HCs surrounded by support cells and mantle cells. HCs afferent innervation is visible in both sections (light stained structures, red *). (**C** and **D**). Higher magnifications of the corresponding areas (blue squares) in successive sections of the wild type and the cnr2 NMs respectively, focusing on the synaptic regions displaying ribbon synapses. (**E** and to **H**) Higher magnifications of each corresponding area (green squares) highlighting two wild type ribbon synapses with tethered vesicles at equidistance that appear highly organized and evenly shaped (white arrowheads). Docked vesicles (white arrows) are also visible (in H) in close vicinity to the PSD (blue arrow). (**F**, **G**, **I** and **J**) Higher magnifications of each corresponding area (orange, red, yellow, and burgundy squares) highlighting 4 different aberrant ribbon synapses in a *cnr2^upr1/upr1^* NM. The ribbons appear bigger and misshapen. The surrounding vesicles are at various distances from the central body, and of different size and shape (magenta arrowheads). Some docked vesicles (magenta arrows in F, I and J) are visible and so is the PSD (blue arrows in F, I and J). Scale bars: in A and B = 5 microns; in C and D = 1 micron; in E to J = 50nm.

Next, we assessed the integrity of ribbons synapses in the retina of wild type and cnr2-KOs larvae. We focused on the outer plexiform layer (OPL) where ribbon synapses in cones pedicles (Figure 6-supplement 1A and D) and rod spherules (not shown) are abundant [67] [23]. The base of the ribbon synapse is anchored at the presynaptic active zone via the arciform density, a retina specific structure where a number of synaptic proteins like Bassoon, RIM2, ubMunc13-2, ERC2/CAST1 are located (for review [68]). These proteins are important for the proper localization of L-type calcium channels (Ca_v_1.4) at the presynaptic plasma membrane. In wild type retina, we repeatedly found ribbons synapses with clearly defined arciform densities (6-Sup 1A and D, blue arrows), opposing equally well-defined electron dense presynaptic plasma membrane (blue arrowheads). The vesicles in the vicinity of the ribbons appeared abundant and evenly-sized (white arrowheads in A and D). However, in mutant retina (B, C, E, and F), most ribbons had incomplete or poorly defined arciform densities (magenta arrows), and the presynaptic plasma membrane appeared blurry and weakly delineated (magenta arrowheads), suggesting disturbed presynaptic active zones. Furthermore, neurotransmitter vesicles in the vicinity of the ribbon synapses appeared scattered, more interspersed, and of variable size (white arrowheads in B, C, E, and F) suggesting that vesicular trafficking was also perturbed. Taken together, the absence of functional cnr2 seemed to affect the ultra-structure of retinal sensory synapses pointing to a putative regulatory role in the eye, reminiscent of our findings in the inner ear and LL.

### Cnr2 affects vesicular trafficking in HCs

To assess the integrity of vesicular trafficking in HCs, we briefly exposed 4dpf wild type and mutant larvae to FM 1-43 live dye and subsequently imaged NMs *in vivo* (L5 in the pLL) from wild type and mutant animals (n= 3/genotype) during 24hour post-treatment (hpt) (Figure 7A, B, and C). At one-hour post-treatment (hpt), the staining did not appear different in wild type and mutant HCs of the NMs (Figure 7. supplement 1, at 1hpt) suggesting that mechanotransduction was intact in mutant HCs. This result was in line with *cnr2^upr1/upr1^* larvae showing no overt behavioral phenotype after sound and vibration stimulation (LC and MB unpublished data).

**Figure 7.**
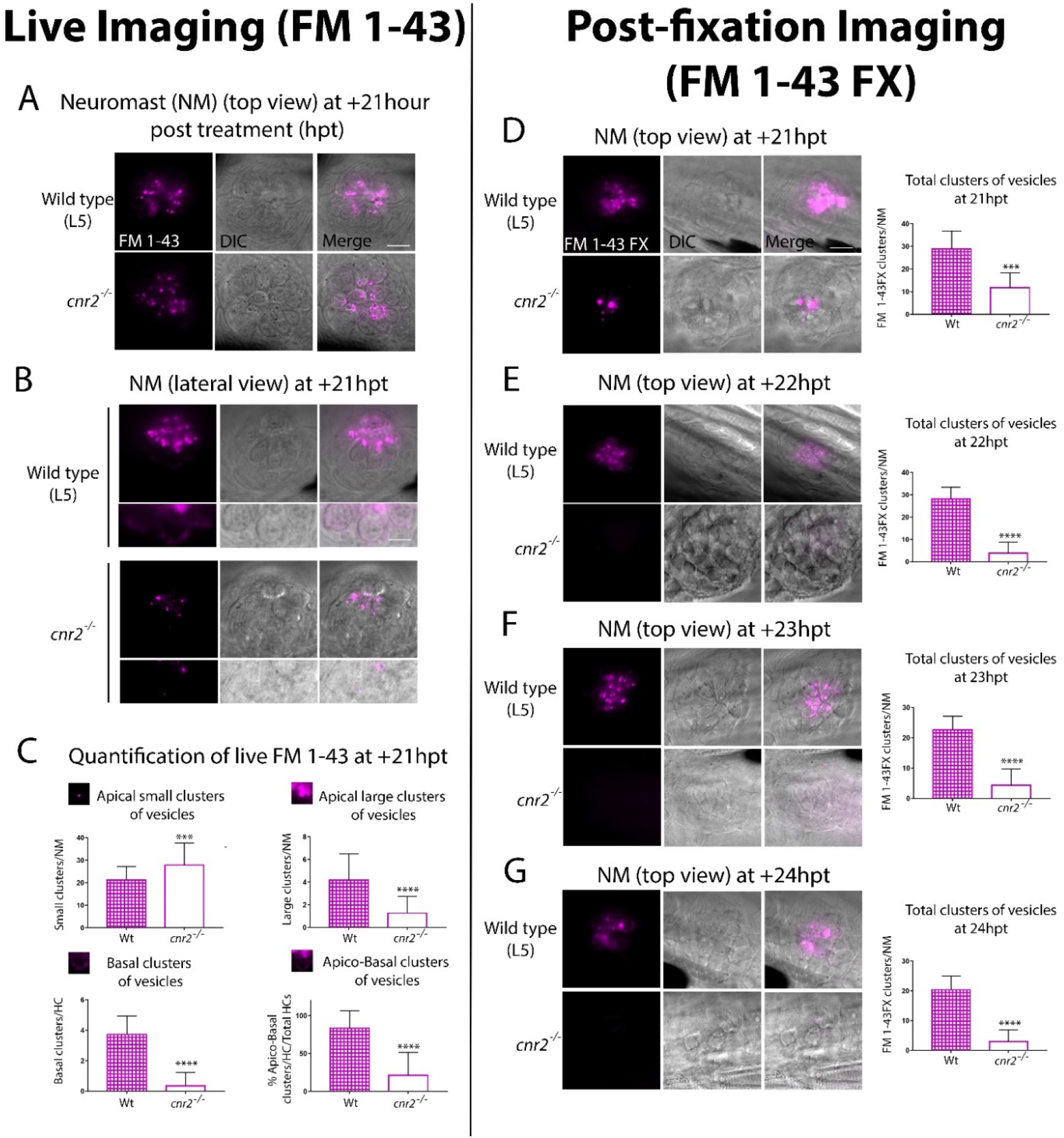
Live (A to C) and post-fixation (D to G) imaging in wild type and mutant NMs after FM 1-43(X) treatment to visualize endocytic vesicular trafficking. (**A**). Top views and (**B**) lateral views of NMs of the pLL (L5) at +21hpt in wild type (top row) and *cnr2^upr1/upr1^* (bottom row). Respective inserts below each panel show the magnified basal compartments of the HCs. (**C**) Quantification of FM-43 residual staining at +21hpt in wild type (checkered bars) and *cnr2^upr1/upr1^* (open bars) of apical small clusters (top left graph), apical large clusters (top right graph), basal clusters (bottom left graph) of vesicles, and % of apical and basal stained HCs /total HCs (bottom right graph). (**D** to **G**). Top views of NMs of the pLL in wild type (top panels) and *cnr2^upr1/upr1^* (bottom panels), showing the vesicular distribution of FM 1-43FX with the respective quantification o number of clusters of vesicles/NM (right graphs), at +21hpt (D), +22hpt (E), +23hpt (F), and +24hpt (G). Scale bars: = 20 microns in A and D, and = 10 microns in B. Significance is represented *** p < 0.003 and **** p < 0.0001.

However, over time and most visible from 18hpt onward, residual FM 1-43 staining was weaker in mutant HCs (Figure 7A, top view at +21hpt). To narrow down on the cellular localization of vesicles, we imaged HCs in lateral views (Figure 7B). In wild type (top panels), we found residual FM 1-43 staining in two clearly distinct cellular compartments, above and below the HC nuclei (visualized in bright field, central and merged right panels). In the apical compartments, we visualized FM 1-43 mostly clustered in vesicular and tubular structures. In the basal compartments, we found a more discreet but very distinct vesicular staining in vicinity to the basal cytoplasmic membrane, possibly coinciding with ribbon synapses location (magnified in bottom half panels). In contrast, mutant HCs (7B, lower panels) showed a much weaker vesicular/tubular apical staining with most clusters appearing much smaller than in wild type NMs. Strikingly, in the basal compartments we found no residual staining (bottom half panels). To quantify the observed differences, we extensively imaged NMs (L1 to L5) in wild type (n = 7) and mutant (n = 8) larvae at + 21hpt (Figure 7C). Using 3D reconstructions, we calculated the percentage of HCs with an apical and basal staining vs. total HCs (= % apico-basal clusters, bottom right graph). We found close to 4 times less apico-basal staining in mutant (open bar) vs. wild type HCs (checkered bar, Wt = 83.65% vs. *cnr2^upr/upr1^* = 21.21% 9, p < 0.0001). Next, we considered apical staining separately which we further partitioned into small (diameter, d > 0.9 μm, top left graph) or large (d > 1 μm, top right graph) clusters of vesicles. Wild type NMs (checkered bars) had ^~^4 times more large apical clusters than *cnr2^upr/urp1^* (open bars) NMs (Wt = 4.171 vs. *cnr2^upr1^* = 1.282; p < 0.0001), but small clusters were predominant in mutant NMs (Wt = 21.29 vs. *cnr2* = 27.9; p = 0.001). The reduced residual apical staining and the increased number of small vesicles in mutant NMs suggested accelerated endosomal turn-over and/or increased apical endocytic activity. Furthermore, in the basal compartments of HCs, we seldomly found residual FM 1-43 in the mutant NMs (bottom left graph: Wt = 3.87 vs. cnr2 = 0.03, p = 0.0001), pointing to a possible alteration of synaptic endo- and exocytosis. Taken together, we observed that vesicular trafficking was severely altered within both apical and basal cell compartments of mutant HCs.

To confirm our findings, we established a time-course starting at + 21hpt (+ 21, +22, +23, and +24hpt) imaging 4dpf fixed larvae (n= 4/ genotype/stage) that we had previously treated with a fixable analog of FM 1-43 (FM 1-43FX, Figure 7D to G). As previously observed, we found a strong reduction of the overall FM 1-43X staining in mutant NMs at +21hpt (top views, 7D left panels). We quantified and averaged the clusters and found a strong decrease in the number of fluorescent clusters/NM (left graph) between wild type (checkered bar) and mutant (open bar) NMs (Wt = 29 vs. *cnr2 ^upr1^* = 11.8; p = 0.0001). At +22hpt (7E) when comparing to + 21hpt, number of clusters in wild type was not significantly different, but dropped drastically in mutant NMs, increasing to a 7-fold difference between genotypes (Wt = 28.2 vs. *cnr2 ^upr1^* = 3.9; p < 0.0001). From this stage onward, in mutant NMs we only found 3 to 4 residual FM 1-43X containing clusters. In stark contrast, in wild type NMs, we observed a slow and steady reduction of stained clusters/NM at +23hpt (7E, Wt=22.63 vs. *cnr2^upr1^* = 4.33; p < 0.0001), and +24hpt (7G, Wt=20.42 vs. *cnr2 ^upr1^* = 3; p < 0.0001), suggesting that this approach was reliably reporting vesicular trafficking/recycling overtime. Taken together, the post-fixation imaging confirmed the strong reduction in residual FM 1-43 live staining that we had previously observed in animals lacking cnr2. Furthermore, the time course revealed a subtle and gradual reduction of FM 1-43X in wild type NMs and highlighted a much more drastic drop in mutant NMs. Taken together, our FM 1-43 experiments demonstrated a strongly altered vesicular trafficking/recycling in HCs of animal lacking cnr2, possibly due to increased endo/exocytic activity in both the apical and basal compartments of HCs. Thus, we postulated that cnr2 function is intimately linked to vesicular trafficking/recycling and mechanotransduction in HCs, ultimately affecting proper LL and auditory function.

### Cnr2 significantly alters swimming behavior in response to sound stimulation in a light dependent manner

Next, we asked if auditory responses were affected in the absence of cnr2. To do so, we monitored swimming behaviors of wild type and mutant animals that we submitted to repeated one-second-long sound stimuli of constant amplitude (S = 450Hz) emitted at 5 min intervals (Figure 8, S1 to S11). To dissociate auditory from visual responses, animals were either first exposed to 90-minute of constant light followed by 90-minute of constant darkness (Fig.8A to E), or vice versa (Fig.8F to J). For animals first exposed to light, as we had previously demonstrated the baseline swimming activity (SA) for mutant animals was lower than for wild type (Fig.8A, yellow box) [8]. In response to sound stimulation (green dashed lines in 8A yellow box, B and D) the SA was only slightly increased in wild type by ^~^14% (black, SA_wt-1_ = 6.80 vs. SA_wt-0_ = 7.65 cm/min, p < 0.0001), but by ^~^63% in mutant larvae (magenta in D, SA_cnr2-1_ = 2.87 vs. SA_cnr2-0_ = 4.67cm/min, p < 0.0001). During the following dark periods upon sound stimulation (green dashed lines in 8A grey box, C and E), the respective increases in SA in wild type and mutant larvae were more similar in absolute values, ^~^100% in wild type (SA_wt-1_ = 2.89 vs. SA_wt-0_ = 5.79 cm/min, p > 0.0001) and ^~^112% in mutant larvae (SA_cnr2-1_ = 1.90 vs. SA_cnr2-0_ = 4.03 cm/min, p < 0.0001), and stronger in wild type in relative values. Taken together, this was suggesting a higher sensitivity of mutant larvae to sound, but mostly when also exposed to light. Interestingly, for animals first exposed to dark, the SA was not significantly different in wild type and mutant larvae neither between, nor in response to sound stimulation in dark (green dashed lines in 8F grey box, G, and I). Thus, this corroborated that mutant larvae sensitivity to sound stimulation was higher when animals were also exposed to light. As expected during the following light periods (Fig.8F yellow box, H and J), mutant travelled less than wild type, but surprisingly more than mutant larvae that had not been previously exposed to sound (compare magenta in yellow boxes in A vs F). This was true before, during, and after sound stimulation when comparing D and J (SA_cnr2-1_ = 4.51 vs. SA_cnr2-1_ = 2. p < 0. 0001, SA_cnr2-0_ = 5.98 vs. SA_cnr2-0_ = 4.67, p < 0. 0001, and SA_cnr2+1_ = 4.52 vs. SA_cnr2+1_ = 2.92 cm/min, p < 0. 0001, respectively), but only for mutant larvae. In wild type larvae with (black in J), or without (black in D) prior sound exposure, the SA was not different before, during, or after sound stimulation (SA_wt-1_ = 6.80 vs. SA_wt-1_ = 6.74 cm/min, SA_wt-0_ = 7.65 vs. SA_wt-0_ = 7.37 cm/min, and SA_wt+1_ = 6.82 vs. SA_wt+1_ = 6.87 cm/min, respectively). Thus, in light periods the SA of mutant larvae with prior exposure to sound stimulation was higher and closer to the wild type SA, which was always unaffected by sound exposure. Taken together, animals lacking cnr2 appeared more sensitive to sound especially when also exposed to light, and less sensitive to light exposure after prior sound exposure. This was suggesting that both auditory and visual sensory systems were affected possibly in an inter-dependent manner in the absence of cnr2.

**Figure 8.**
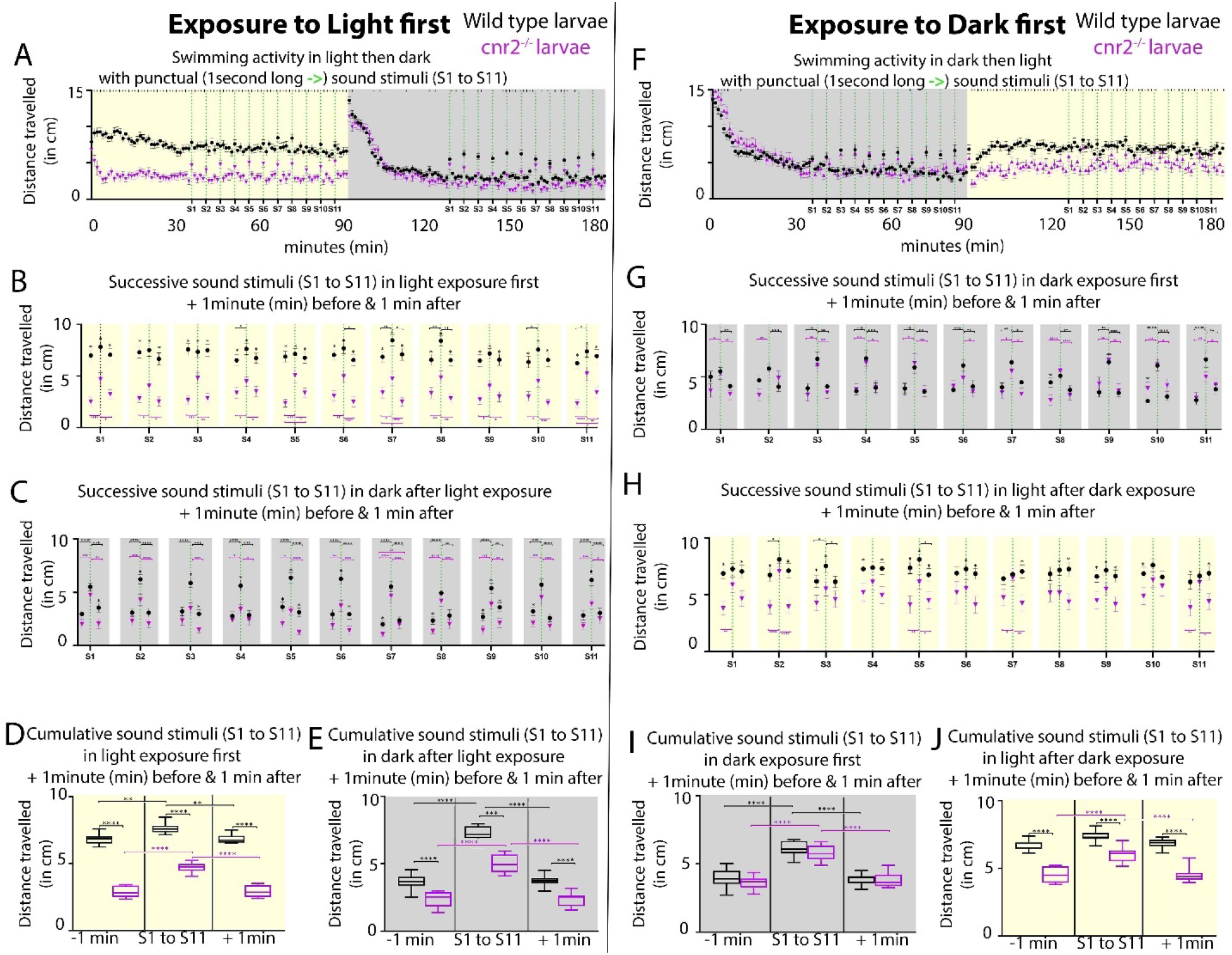
Swimming activity recorded in 6-day post-fertilization (dpf) wild type and *cnr2^upr1/upr1^* larvae after punctual sound stimuli (S1 to S11) during light or dark exposure. (**A**). Averaged swimming distances travelled /larva/min by wild type (black dots) or mutant (magenta triangle) animals, that were first submitted to a 90-minute light period (yellow box = maximum and constant intensity of 385Lux), followed by a 90-minute dark period. At minute 36, larvae were exposed to a one-second-long sound stimulus (S, vertical green dashed line at 450Hz) and then repeatedly at 5min intervals (S1 to S11, yellow box). At S12, light was turned off and animals were left unstimulated for acclimation to dark for 35 min (grey box). At minute 126, sound stimulations were resumed (S1-S11, green dashed lines in grey box). (**B** and **C**) Averaged swimming responses in light (B, yellow box) and dark (C, grey box) periods during all successive sound stimulations (S1 to S11) showing the average distance travelled during the one-minute preceding, including, and following each individual sound stimulus (S1 to S11, green dashed lines). (**D** and **E**) Averaged swimming distances in light (D, yellow box) and dark (E, grey box) periods showing the compiled results for all one-minute preceding (−1min), one-minute including (S1 to S11), and one-minute following (+1min) the 11 sound stimulations. (**F** to **J**). Same modalities as in (A to E) except that animals are now first exposed to dark for a 90-minute followed by a 90-minute light period. Error bars represent the standard errors of the mean (SEM) and significance is indicated by * (* = p < 0.05, ** = p < 0.01*** p < 0.001 and **** p < 0.0001, ns. were omitted for clarity), comparing wild type to mutant swimming activity, or as indicated by the color coordinated parentheses.

Based on the perturbation that we observed in vesicle distribution at the ribbons in mutant retina, we hypothesized that mutant animals challenged by even slight variations in light intensity would not be able to fully readjust during gradual changes. To test this, we first reproduced our previously published experimental setup in which we recorded wild type and mutant larvae SA after an initial 30min adaptation to dark, followed by 4 successive 10-minute light/10minute dark periods (Figure 8-supplement 1A [8]). As expected, in mutant larvae (magenta squares) the SA was significantly reduced in light and increased in dark periods. Notably the difference with wild type SA (black dots) was not significant at the end of light and beginning of dark periods, suggesting that mutant larvae were slower to adapt to light, but also less challenged in dark. Next, instead of abrupt changes to full light or complete darkness, we progressively increased in one-minute increment to full light (100%) and then decreased to full darkness (0%) over the course of a 10min period, for 4 successive cycles (Figure 8-supplement 1B). Responses from one cycle to the next were highly consistent and showed that the swimming behavior was significantly different at all recorded time points between wild type and mutant animals, for the exception of the minute of, and the minute immediately after full darkness. Mutant larvae swum significantly less at all other time points, implicating cnr2 function in detection of even slight light changes. Interestingly, the global trends in SA over all 4 cycles was identical in wild type and mutant larvae, but at a much lower activity level in mutant larvae (compare black and magenta tracks/slopes in each cycle, SA_WT_ ^~^ 8cm/min vs. SA_cnr2_ ^~^ 5cm/min). This suggested that adaption to light changes was occurring in mutant larvae, but less effectively, thus corroborating findings in adult CB2-KOs mice [69]. Taken together, this suggests a conserved role for cnr2 in adaptation to light that appears early during development and is maintained into adulthood.

## DISCUSSION

### Cnr2 expression in physiologically active and mature HCs of the Inner Ear and the LL

We demonstrated strong *cnr2* transcriptional expression starting at 3dpf in zebrafish larva in the mechanoreceptors or hair cells (HCs) of sensory epithelia (SE) of the inner ear and lateral line (LL). We further showed that the early development of those SE and the overall morphology of those organs was unaffected in *cnr2^upr1/upr1^* animals which are totally lacking cnr2 [8]. Thus, this was suggesting that cnr2 was not involve in early development of either organ, but rather had a role in maturation and/or physiology of HCs.

Little is known about the expression or function of either cannabinoid receptors (CNR1 or CNR2) in the ear and hearing. Audiograms and measurements of gap detection thresholds in Cnr1-KOs mice showed impaired hearing abilities at higher frequencies but enhanced gap detection thresholds [70]. Those differences were attributed to Cnr1 function in the auditory brainstem where it is highly expressed [71–73]. For Cnr2, reports demonstrate the involvement in inflammation responses in the outer [74] and middle ear [75]. In the inner ear, a recent report documented spontaneous Cnr2 expression in the cochlear canal of adult rats [11]. Strong immunolabelling of Cnr2 was found in the *stria vascularis* (SV), and maybe more surprisingly in all inner HCs (IHCs), as well as in the afferent neurites and cell bodies of spiral neurons [11]. All these localizations were confirmed independently, and additional cochlear expression was found in the spiral ligament (SL), the outer HCs (OHCs) and a subset of support cells (SCs), namely the inner and outer pillar cells (IPCs and OPCs respectively) [12]. Furthermore, this group showed that Cnr2 was co-localizing with markers of fibroblasts in the SL, basal cells in the SV, and ribbon synapse (Ribeye B or CtBP2) in the IHCs [12].

The underlying mode of action of Cnr2 in any of those cells/tissue remains to be determined. However, both groups demonstrated that Cnr2 expression was up-regulated after Cisplatin exposure [11, 12]. This was strengthening an earlier claim for an anti-apoptotic role for Cnr2 in cisplatin-treated cultured cells of an auditory cells line [76]. In rats, intra tympanic pretreatments with Cnr2 agonist or antagonist (JWH015 and AM630 respectively) before cisplatin application, modulated cochlear inflammation, but pointed to a protective role of Cnr2 against several other ototoxic effects. First, it prevented apoptosis in OHCs. Second, it reduced loss of ribbon synapses in IHCs, and third, it maintained Na^+^/ and K^+^-ATPase activity in SV and SL [12]. All three aspects are strong contributors to optimal hearing. Furthermore, *in vivo* Cnr2-Knockdown (KD) showed that at least at lower frequencies, Cnr2 could mitigate cisplatin-induced hearing loss [12], thus strengthening the hypothesis of a protective role for Cnr2. Interestingly, presbycusis, or age related hearing loss, is not only linked to HC loss, but also to SV degeneration with reduction of Na^+^ and K^+^-ATPase activity, the latter ultimately resulting in an energy starved cochlear amplificatory system [77, 78].

Similarly, a cardioprotective role for Cnr2 activation was described [79], but it remains to be demonstrated if Cnr2 can offer protection against all forms of hearing losses. Furthermore, future therapeutic approaches will need to be Cnr2 specific and topical, because at least for Tinnitus (intermittent or constant phantom noise that has been mostly linked to auditory brainstem defects), cannabinoid treatments had negative effects [73, 80, 81].

### Cnr2 dependent maturation of ribbon synapses

We found strong alteration in the sensory synapses of HCs of the LL in the pre- and post-synaptic elements, as well as in their alignment in 5dpf *cnr2^upr1/upr1^* animals. Ribbon synapses formation and maturation has been well characterized in this very same context and at the developmental stages that we assessed (for review [82]). We found that all wild type animals exhibited the previously described characteristic mature synapses in all HCs of NMs in the anterior LL (aLL), which develops first, as well as in the most rostral NMs of the posterior LL (pLL). As expected, we found more immature ribbon synapses in the most caudal wild type NMs which are constantly added as the animal is growing [83]. In stark contrast in HCs of mutant animals, the NMs from the aLL and rostral pLL had mostly immature or even grossly misshapen ribbon synapses. Differences became less striking following an antero-posterior gradient in the pLL and were not always significant in the tail. Taken together, this was pointing to an involvement of cnr2 in the maturation of the sensory synapse.

Developmental maturation of ribbon synapses was previously extensively described in HCs [84], but has never before been linked to cnr2 function. The first mature and functional HCs can be found in the LL and inner ear as early as 3dpf, even if the maturation process is ongoing in both structures into adulthood [13, 85]. HC innervation also happens early and is extensively documented in the LL [86–88], and it was demonstrated that innervation regulates ribbon synapses development and maturation [32]. Interestingly, we did not find overt defects in the sensory or motor innervation of NMs, thus suggesting that the defects observed in the sensory synapses were HC autonomous, but this remains to be tested. However, we systematically found altered Maguk staining in the post-synaptic elements. Instead of being focalized in close vicinity to Ribeye b and aligned with it as we found, and was previously described in wild type animals, it appeared diffuse and generally weaker in the mutant animals. Notably, we found similar alterations in the inner ear. The importance of physiological activity for maturation of the ribbon synapse is well known and has gained a lot of attention in these last years (for review [89]) especially in regard to synaptopathies (discussed below). Our work clearly pointed to a cnr2-mediated cross talk at the afferent synapse, thus offering a novel mechanistical link between maturation and activity.

### Cnr2 modulation of synapse plasticity

We found a perturbed distribution of voltage gated L-type Ca^+^ channels (Ca_v_1.3) at the active zone in *cnr2^upr1/upr1^* animals. The need for tight regulation of Ca^2+^ exchange at the sensory synapse had been extensively demonstrated in HCs of mammals [37, 90, 91] but also Fish [28, 38, 39]. An intimate relationship between ribbon size and Ca_v_1.3 channels density and distribution was repeatedly highlighted [39–41]. In physiological situations bigger ribbons were shown to have more associated Ca_v_1.3 channels and larger calcium signals [92, 93], but it was unclear if this ultimately translated into higher afferent activity. Additional work showed that when post-synaptic elements were unaffected and ribbon synapses’ size increased, clustering of Ca_v_1.3 channels was reduced at the active zone [31], but both global and ribbon-localized calcium signals were increased, suggesting an alternate mode of Ca^2+^ levels regulation.

In conventional synapses, the endocannabinoid system (ECs) governs synaptic plasticity via a well described retrograde signaling that will result in short- or long-term depression of the presynaptic element (for review [2] [94]). Short-term plasticity mostly involves direct G protein-dependent inhibition of Ca^2+^ influx through voltage-gated Ca^2+^ channels (VGCCs) [95–97]. CNR1 activation was shown to exert feedback inhibition of N, P/Q -type VGCCs in the CNS [98–100], but also of L-type channels in smooth muscles [101] and bipolar neurons [102]. Furthermore, accumulating evidence demonstrated that CNR2 can also modulate synaptic activity in a variety of neurons, presumably through identical mechanisms [103–105]. Thus, in HCs one possibility is that Cnr2 regulates mechanotransduction by modulating either global or local Ca^2+^ currents. This remains to be assessed and could be done by *in vivo* electrophysiology recording of HCs of the LL and inner ear using techniques which have been specifically developed in both organs in the developing and adult zebrafish [106–108]. Measurements of whole cell and local Ca^2+^ currents in HCs of *cnr2^upr1/upr1^* animals could be further coupled to calcium imaging using some of the existing transgenic lines [109]. Finally, concomitant to Ca^2+^ influx, activation of K^+^ efflux is also directly controlled by ECs signaling [98, 110]. Interestingly, recent work highlighted how only a subset of HCs in each NM were active while others were silent due to an unknown regulatory mechanism linked to K^+^ levels [111]. Our discovery of a modulatory role of cnr2 at the HC ribbon synapse offers a tantalizing mechanism that begs testing.

ECs-dependent long-term synaptic plasticity involves inhibition of adenylyl cyclase and downregulation of the cAMP/PKA pathway which will ultimately result in inhibition of neurotransmitter release [112, 113]. We demonstrated that vesicles at the mutant ribbon synapse were uneven in size and shape, and that the tethering to the ribbon was irregular. The ribbon tethered vesicles represent the ready-to-release pool (RRP) available to support continuous transmission and constitute a timing system for delivering those vesicles to the plasma membrane in a synchronized manner (for review[21] [24, 27, 35]). Electron tomographic reconstructions of inhibited or stimulated HCs showed that depolarization was creating a gradient in size of the vesicles’ size [114], raising the possibility that the observed phenotype in the mutant HCs resulted from defective inhibition or regulation by Cnr2. This could be readily tested by capacitance measurements to establish if vesicular fusion is affected. HC synapse function can be further measured by electrophysiological recording from afferents which has been perfected in HCs of the LL [31, 115, 116].

We next showed that neurotransmitter vesicles in the vicinity of the ribbon synapse were affected in size, shape, and number at the active zone in cnr2 homozygotes. The mode of vesicle release is still a matter of debate in the field with evidence of both uniquantal release (UQR) and coordinated multiquantal release (MQR) in which coordinated synaptic vesicle (SV) exocytosis occurs at the ribbon active zone (AZ) (reviewed in [35]). However, what is agreed on is the necessity of appropriate coupling of exo- and endocytosis to allow the indefatigable synaptic transmission at ribbon synapses [35]. Endocytosis is taking place in the peri-active zone which was demonstrated in the photoreceptor synapses [117], and in the frog saccular HCs [114]. Three endocytic mechanisms have been described (1) ultra-fast clathrin independent, (2) fast bulk endocytosis [118], and (3) slow clathrin-mediated [119]. Cnr2 could potentially modulate any or all those steps. Crossing the cnr2 mutants with loss-of-function mutant lines in proteins that have been involved in those mechanisms specific to the ribbon synapses like otoferlin will be helpful to address that (for review [27]).

Finally, we showed that the overall vesicular trafficking seemed accelerated in mutant HCs which unlike the wild type larvae, had very little residual staining in the apical and basal compartments from 24hour post staining (hpt) onward. Notably, the HC specific apical endocytosis appeared unchanged, thus suggesting a difference in exo-endocytosis which could be restricted to the ribbon synapses but might also be due in part to constitutive membrane trafficking in other cell parts. To distinguish between those possibilities, a recently developed technique which is coupling FM 1-43 staining to photo-oxidation will allow to focus on synaptic trafficking only [65].

A less well explored mode of action for ECs was demonstrated for CNR1 which was found anchored in the external mitochondrial membrane, where it directed cellular respiration and energy production in murine neurons, and participated in regulation of retrograde inhibition [120]. Recent work in HCs of the LL highlighted how the ribbon size was modulated by mitochondria and how mitochondrial Ca^2+^ was participating in synaptic function [121]. It remains to be demonstrated if Cnr2 is expressed in mitochondria in the vicinity of the ribbon synapse. If so, a provocative hypothesis to explore will be that mito-Cnr2 exerts a role like mito-Cnr1 in neurons, but specifically in sensory cells.

### Behavioral changes in cnr2 mutant in response to sound stimulation and light changes

When testing the swimming behavioral of animals lacking cnr2, we found that homozygote *cnr2^upr1/upr1^* larvae appeared more sensitive to sound especially when also exposed to light. Hyperacusis is a hyper-sensitivity to sound that was found associated with noise-induced damaged ribbon synapses in a mouse model [122]. Synaptopathies get often undetected by audiograms because patients have normal hearing thresholds, but reduced supra-thresholds and it has been called the hidden hearing loss [123]. Additional unexpected finding in aged mice that had been young-exposed to noise at levels that produced only moderate threshold shift and no HCs loss, was that acute loss of synapses and peripheral terminal of the spiral neurons was presaging later ganglion cells losses [124]. This was suggesting that defective sensory synapses were setting the stage for neurodegeneration. This raises the possibility that we might be able to observe acute HCs degeneration in homozygote *cnr2^upr1/upr1^* adults. However, HCs regenerate in fish unlike in mammals, and our earlier observation was that HC regeneration was unaffected in the larval LL. It will be of interest to verify if this remains true in the adult inner ear and LL.

Notably, mutant animals were responding differently to sound but more so when in light. Pioneer work has recently shown a similar relationship between light and noise sensitivity in mice, demonstrating a higher sensitivity to noise in dark and showing the presence of a molecular clock in the cochlea, linking it to the circadian rhythm [125]. Unlike mice, zebrafish are diurnal which might explained the greater sensitivity to noise in light rather than in dark. The presence of a molecular clock in fish HCs as well as its hypothetical modulation by cnr2 will be an interesting avenue to pursue. Interestingly, pinealocytes in the pineal gland, a major player in the control of circadian rhythms have ribbon synapses that presented an altered morphology in mutant larvae, (LCC and MB unpublished), which warrants further exploration.

We had previously shown that *cnr2^upr1/upr1^* larvae were behaving differently in light and dark [8] which we reproduced here, and we further showed that mutant larvae were slower to adapt to light changes. Anecdotal reports of improved night vision after cannabis ingestion [126, 127] and glare recovery impairment experiments [128] have mostly linked altered vision to CNR1. However, the retina of adult mice showed no alteration neither in Cnr1-KOs nor Cnr2-KOs, and only the latter required more adaptation time to light [69]. Cnr1 protein expression in adult retina was extensively described in several cell types and in various species (for review [9]) including goldfish [10]. In the developing retina, expression of Cnr1 along with other components of the endocannabinoid signaling pathway [129, 130] were described in embryonic [131] and postnatal rat [132], as well as embryonic chick [133]. Pharmacological manipulations with Cnr1 and Cnr2 agonists and inverse agonists suggested an involvement of ECs in the retinothalamic development which needs further investigation [134]. However, the expression of Cnr2 in the adult retina remains controversial because of debated specificity of available Abs [135, 136] and has not been explored in the developing retina, which needs to be addressed.

We found subtle but consistent alterations in the retina of mutant *cnr2^upr1/upr1^* larvae. Most ribbons had incomplete or poorly defined arciform densities where the base of the ribbon is anchored. The main component of this electron dense ultra-structure is Bassoon, which when functionally disrupted results in free floating ribbons [137]. The active zone had a blurry and weakly delineated appearance and the surrounding vesicles appeared scattered, interspersed, and of variable size. Taken together, this was suggesting perturbation of the vesicular trafficking in the presynaptic active zone that needs further investigation. Ribbons in photoreceptors are highly dynamic and their size is well known to vary with illumination [33] and diurnal signals in mice and fish [68, 138–142]. Behavioral experiments allowing to discriminate between the Cnr2 exerted regulation of individual sensory systems (pineal gland, retina, inner ear, and LL) will be highly informative. The deciphering of each relative contribution will be paramount in providing a holistic understanding of the Cnr2 role(s) in the regulation of individual and combined sensory inputs.

### Cnr2 activation: a potential therapeutic route for Synaptopathies

We have established a clearl link between cnr2 and the sensory synapse. Auditory or cochlear synaptopathies are a specific type of sensory hearing loss in which HCs appear intact and can detect sound stimulation, but are unable to transmit the signal at the sensory synapse [143]. The origin can be genetic like loss-of function mutations in the *Vesicular glutamate transporter-3 gene (VGLUT3*) causing progressive non syndromic hearing loss [144], that was independently identified in zebrafish [145] and in mice [146]. Zebrafish HCs are remarkably similar to mammalian HCs [147] and there is a strong gene conservation from fish to mammals [148]. Thus, a growing number of mutant lines are providing excellent tools for hearing disorder modeling [85, 149]. More recently it became evident that damage to ribbon synapses represent a highly prevalent form of acquired sensory hearing loss [150, 151], mainly caused by aging [152] and noise-induced damage [124, 153]. Aging as well as overexposure to noise, both result in dramatic swelling of the afferent dendrites at the ribbon synapse which can be prevented by pretreatment with AMPA/Kainate antagonists of the post-synaptic glutamate receptors as well as by pharmacological blockage of glutamate release, suggesting glutamate excitotoxicity [154]. Notably, sensitivity to aminoglycosides might also be related to damage in ribbon synapses as a primary effect which will eventually be followed by HCs death depending on the dose [155]. Furthermore, and as discussed above cisplatin ototoxicity was affecting ribbons synapses and was mitigated by Cnr2 pharmacological manipulations [12], thus offering a promising therapeutic approach to correct or prevent synaptopathies.

## CONCLUSIONS

We showed for the first time a clear relationship between the maturation and function of the ribbon synapse (RS) and the endocannabinoid system (ECs) in two sensory systems of a developing vertebrate. Our work explored how a loss-of-function mutation in the *cnr2* gene was linked to defective swimming responses triggered by sound and light in zebrafish mutant larvae. First, we demonstrated for the first time the expression of *cnr2* in HCs of the LL and the sensory patches of the inner ear in larval zebrafish, which was concordant with previous expression studies in adult rodents and derived auditory cell lines [11, 12, 76]. Second, we showed strong perturbations in several components of the sensory synapse in the mutant *cnr2^upr1/upr1^* larvae which became increasingly obvious as HCs were maturing. We noted similar perturbations in the RS in the mutant retina. Third, we linked these morphologic alterations of the RS to an altered cellular physiology by showing that the vesicular trafficking in HCs was strongly perturbed. Fourth, we illustrated differences in swimming activity in response to sound or light stimulation in mutant *cnr2^upr1/upr1^* larvae, therefore, underlining the relevance of the observed phenotypic differences. Taken together, we presented alterations linked to the absence of cnr2 in developing zebrafish larva at the ultra-structural, structural, cellular, and physiological levels and ultimately linked them to behavior. Our work strongly suggested a pivotal role for CNR2 in the regulation of mechanotransduction in HCs. Furthermore, our data implied that the CNR2 mediated regulation might be common to other sensory systems.

## MATERIELS & METHODS

### Key Resources Table

Key Resources Table:
| Reagent type, species or resource | Designation | Source/reference | Identifier/Dilution |
| --- | --- | --- | --- |
| Genetic reagent (*Danio rerio*) | Wild Type | UPR bred fish stock | TAB-5 (Tubingen x AB) |
| Genetic reagent (*Danio rerio*) | Wild Type | Gift from Shawn Burgess 'Lab | NHGRI-1 |
| Genetic reagent (*Danio rerio*) | cnr2-KO | Acevedo, Colón & al., 2019 | *cnr2<sup>upr1/upr1</sup>* |
| Gene (*Danio rerio*) | *cnr2* | Ensembl | ENS DARG00000039970.7 |
| Commercial assay or kit | RNA Clean & Concentrator | Zymo Research | R1016 |
| Commercial assay or kit | DNA Clean & Concentrator | Zymo Research | RD4014 |
| Commercial assay or kit | SuperScript III One-Step RT PCR Kit | Sigma Aldrich | 12574-026 |
| Chemical compound | FM 1-43 | Invitrogen | T35336 |
| Chemical compound | FM 1-43 FX | Invitrogen | F35355 |
| Antibody | Ribeye b (Mouse Monoclonal) | Gift from Teresa Nicolson's Lab | d = 1/2000 |
| Antibody | Ribeye b (Rabbit Polyclonal) | Gift from Teresa Nicolson's Lab | d = 1/500 |
| Antibody | Ca<sub>v</sub>1.3a (Rabbit Polyclonal) | Gift from Teresa Nicolson's Lab | d = 1/1000 |
| Antibody | Anti-Maguk/(k28/86) (Mouse Monoclonal) | NeuroMab, Davis | AB10673115<br>d= 1/1000 |
| Antibody | Znp1 (Mouse Monoclonal) | Hybridoma Bank | AB2315626<br>d = 1/100 |
| Antibody | Zn12 (Mouse Monoclonal) | Hybridoma Bank | AB531908<br>d= 1/100 |
| Antibody | Myosin VII (Mouse Monoclonal) |  | #25-6790<br>d= 1/100 |
| Antibody | II Ab- Rabbit Alexa-Fluor 488 | Thermo Fisher | A11034<br>d= 1/1000 |
| Antibody | II Ab- Mouse Alexa-Fluor 568 | Thermo Fisher | A11031<br>d= 1/1000 |
| Machine/Equipment | Zebrabox<sup>®</sup> | Viewpoint, France |  |
| Software, algorithm | GraphPad Prism | GraphPad Software | www.graphpad.com |
| Software, algorithm | Zen Lite 2.6 | Zeiss | www.zeiss.com |
| Software, algorithm | Zen Blue | Zeiss | www.zeiss.com |
| Software, algorithm | FIJI | PMID: 22743772 |  |
| Software, algorithm | ZebraLab | Viewpoint, France |  |
| Microscope | Zeiss Axio Imager Z2<br>Coupled to Confocal<br>Laser Scanning<br>Microscope (LSM800) | Zeiss |  |
| Microscope | Inverted Axiovert | Zeiss |  |

### Ethical statement

We carried-out experiments in accordance with the guidelines and protocols approved by the IACUC (#A880110) of the University of Puerto Rico - Medical Sciences Campus (UPR-MSC).

### Zebrafish care and husbandry

We performed animal care and husbandry following previously published protocols [156] and NIH guidelines. We used zebrafish (*Danio rerio*) for all experiments, which we raised and maintained in the UPR-MSC Satellite Fish Room facility according to standard procedures as recommended [156]. We raised and kept all fish at 28°C on 14:10 hour light/dark cycles on a recirculating system (Techniplast^®^). Water supplied to the system was filtered by reverse osmosis (Siemens) and maintained at neutral pH (7.0–7.5) and stable conductivity (1,000 μS/cm) by adding sea salt (Instant Ocean^®^). This water is referred to as system water (SW). After each cross, we collected, rinsed, and raised embryos for the first 24-hour post fertilization (hpf) in SW with methylene blue (0.2 %). After 24-hpf, we raised fertilized and anatomically normal embryos in SW at 28 °C on 14:10-hour light/dark cycles until 6-day post fertilization (dpf). We only used larvae devoid of anatomical abnormalities and exhibiting upright swimming for further experiments. We did all developmental staging according to [157].

### Zebrafish lines

We bred, maintained, and staged wild type (TAB-5 = Tubingen x AB, and NHGRI-1) and *cnr2^upr1/upr1^* mutants as previously described [8]. To genotype fish, we fin-clipped them once they reached adulthood (=3 months old) and digested fins in 30μL 50mM NaOH (Sigma Aldrich) at 95°C for 20 minutes (min). Next, we added 30μL 100mM Tris-HCl, and PCR-amplified fragments from the *cnr2* target region using the following primers (Forward: 5’-GACCACACAAGAGCAGAAAGC-3’, and Reverse: 5’-GACGATCCAACCAGGTTTTG-3’) as stated previously [8].

### Whole-Mount *In-Situ* Hybridization (WISH)

We extracted total RNAs using trizol (TRI-Reagent, Sigma Aldrich) from 5dpf larvae to synthetize sense and antisense probes. We used the SuperScript III One-Step retro-transcription kit (Sigma 12574-026) and designed gene specific primers (GSPs) for *cnr2* (Forward: GATCAAGAAGCTACGACTGTGC, and Reverse: ACTACCACTCACTGCCGGAT) with T7 (atgctaatacgactcactatagggaga) and T3 (atgcattaaccctcactaaaggga) promoter sequences attached to the forward and reverse primers, respectively. The expected amplicon length was 1,080bp. We performed WISH as described previously with minor modifications [158]. First, we dechorionated embryos and rehydrated in methanol/10X PBS gradient solutions (75% methanol, 50% methanol, 25% methanol) for 5 minutes (min) and then rinsed in PBST (1%Tween 20) for 5min. We bleached animals older than 24hpf with 30% H_2_O_2_ (Sigma) for approximately 20min. Next, we digested larvae older than 2dpf with 2mg/ml of Proteinase K (Ambion) for 7min, and later fixed at room temperature (RT) (4% paraformaldehyde) for 45min. After rinsing 5×5min in Blocking Buffer for WISH (1X P880110BS, 0.1% Tween 20, 0.1% BSA, 1% DMSO), we pre-hybridized larvae in hybridization mix (HM) (50% Formamide, 5X SSC, 1mg/mL Yeast RNA, 50μm /mL Heparin, 0.1% Tween 20, 5mM EDTA, 9mM Citric Acid in DEPC treated water) for 4-6 hours. Next, we hybridized all specimens with the pre-heated antisense probe (final concentration = 1ng/μl) overnight (O/N) at 65°C. We washed larvae in a series of HM solutions in 2X SSC (75%, 50%, 25%) for 10min each, followed by two 30min washes of 0.2X SSC, all at 65°C. Next, we rinsed all animals in 0.2X SSC gradient solutions in PBST (75%, 50%, 25%) for 10min each at RT. We pre-incubated all specimens in Blocking Buffer for 4-6 hours and then incubated in pre-absorbed anti-DIG-AB against fish powder in blocking buffer (1/3000) O/N. Finally, we rinsed all animals in PBST 6×15min followed by two 5min washes in Alkaline Phosphatase Buffer (APB) (100mM Tris pH9.5, 50mM MgCl2, 100mM NaCl, 0.1% Tween 20, levamisole). Revelation was performed using BM Purple (100%) in the dark.

### Immunohistochemistry (IHC)

We fixed animals O/N (PFA 4%) and washed 3×5min in PBST (PBS 1X, 0.1% Tween 20). Next, we stored at −20°C in methanol (100%). When ready to perform IHC, we rehydrated specimens using serial dilutions (MeOH 75%/PBST 25%, MeOH 50%/PBST 50%, MeOH 25%/ PBST 75%, PBST 100%) for 10min each. We then replaced PBST with cold Acetone for 7min at −20°C, and then rehydrated in PBST with 3×5min washes. We digested larvae with 1mg/mL of collagenase in Blocking Buffer (PBST, 10% Goat Serum, 0.1% BSA) for 35min. Next, we washed all specimens with Blocking Buffer 5×5min, after which we pre-incubated them in fresh Blocking Buffer for 4-6 hours followed by primary antibody incubation O/N. Next, we rinsed larvae in PBST 4×15min and pre-incubated them in Blocking Buffer for 4-6 hours before incubating O/N in secondary antibody. Next, we rinsed larvae 4×15min in PBST and mounted them for imaging in poly-Aquamount (PolySciences). The following affinity-purified primary antibodies were a kind gift from Dr. Teresa Nicolson [39] and generated against *Danio rerio*: mouse monoclonal against Ribeye b (amino acids 12-33; Open Biosystems, Huntsville, AL) and rabbit polyclonal against Ribeye a (amino acids 1-466; Proteintech, Chicago, IL, USA), Ribeye b (amino acids 4-483; Proteintech) and Ca_V_1.3a (amino acids 42-56; Open Biosystems). We used the K28/86 (NeuroMab, Davis, CA, USA) to label MAGUKs. Secondary antibodies used were Alexa-Fluor 488 Anti-Rabbit and Alexa Fluor 568 Anti-Mouse.

### FM 1-43 and 1-43X Hair cell (HC) staining and live imaging

We used FM 1-43 live dye (Invitrogen, #T35356) and its fixable analog (FM 1-32FX, Invitrogen, #F35355). For either, we exposed larvae for 30 seconds (sec) at the desired stage to 3 μM FM 1-43(X), rinsed 3 x 30sec in SW. For the fixable version and at the desired time point post-treatment we anesthetized in ice cold water and subsequently fixed animals (PFA 4%, O/N). Next day we washed larvae 3 x 5min in SW before mounting them on slides (see above in ICH).

For live imaging, we mounted larvae at the desired post treatment time point using 2% low-melting agarose (LMA) in bottom coverslip chambers. Imaging was performed on an inverted Axiovert (Zeiss) creating z-stacks (in 1μm steps) from an identified NM in the pLL (L5) using the 63X DIC oil-immersion objective. The first stack was recorded as soon as animals were mounted in LMA, and each following stack was recorded in 30min intervals for a period of 24 hours. We recorded the first 4-hours immediately post FM 1-43 staining. We used these z-stacks and time-point recordings to re-construct a 24-hour post-treatment timeframe.

### Semi-thin and ultra-thin sections for Transmission electron microscopy (TEM)

Larvae were prepared as described previously [159]. Briefly, larvae were fixed overnight in 2.5% glutaraldehyde (Sigma) and 4% paraformaldehyde prepared from paraformaldehyde (Sigma) in 0.1M sodium cacodylate buffer (Sigma). Larvae were then rinsed and post-fixed 1h at room temperature in reduced osmium (1:1 mixture of 2% aqueous potassium ferrocyanide) as described previously [59]. After post-fixation, the cells were dehydrated in ethanol and processed for Epon (Sigma) embedding. Semi-thin sections (300 nm) were cut and collected on a glass slide, and subsequently stained using toluidine blue (Sigma). The analysis and imaging were done on an inverted Zeiss Axiovert200M. Ultra-thin sections (80 nm) were cut on a Reichter-E ultramicrotome, collected on copper grids and stained with lead citrate (Sigma) for 2 min. Sections were then examined with a CM 10 Philips electron microscope at 80kV. We performed serial sectioning to follow the same ribbon in three to five sections on the same grid.

### Confocal microscopy

For confocal imaging we processed larvae as previously described [159]. Briefly, IHC processed larvae were mounted in poly-Aquamount (Polysciences) on slides prepared with fenestrated tape in 3 layers to avoid squashing them. Acquisitions were performed on a Zeiss Axio-Imager Z2 coupled to a confocal laser scanning microscope (LSM800) with a Pan-Apochromat 63×/1.40 oil-immersion objective. It is to be noted that staining in the mutants were often considerably weaker and when keeping the same setting used for wild type NMs and ears, we would often have no signal at all making the comparison impossible. Therefore, we adjusted the laser setting as necessary to obtain an optimal image for each NM. When comparing we would integrate the acquisition differences in our final assessment.

### Image Acquisition and Post-Processing

For FM 1-43 staining, we acquired images on an inverted Axiovert (Zeiss) with Axiovision and post-processed as needed with the Zen Lite 2.6 software. Confocal images were acquired as described above with maximal projections of z-stacks created and analyzed using Zen Blue software. Final figures were created with Adobe Photoshop and Illustrator.

### Swimming activity tracking

To measure swimming activity, we conducted behavioral assays using a previously described setup [160]. In brief, 24hour prior to the experiment, we plated one individual larva per well in a 48-well plate (Greiner bio one, CELLSTAR^®^) with 450 μl of SW. The next day and prior to the experiment we topped up each well to 500 μl of SW and performed a health check. Only healthy (with inflated swim bladder, no obvious external damage and swimming upright) were used for the behavioral testing. We placed the 48-well plate into the Zebrabox^®^ (Viewpoint, France), which is an isolated recording device with a top camera and infrared light-emitting base where we controlled temperature, light intensity, and sound/vibration stimulations. We recorded swimming activity 1minute per minute using the Viewpoint tracking software (ZebraLab).

### The photo-dependent response (PDR) and the gradual-PDR (g-PDR)

In the photo-dependent response (PDR) as developed in [160], we first habituated larvae for 30 minutes (adaptation/incubation) in the dark (I_min_= 0 Lux = 0%) followed by a sharpen transition to light at maximum intensity (I_max_= 385 Lux = 100%) for 10 min followed by 10min of dark. We proceeded with four successive cycles (80 minutes) for a total recorded time of 110 minutes.

In the gradual-PDR (g-PDR), we modified the luminosity intensity gradient minute per minute to reach the maximum (I_max_) or minimum (I_min_) light intensity at the end of each 10-minute interval ramping from 0-100% in light and back to dark from 100-0%. Specifically, we changed the light intensity by a factor of +/- 10% (= 38.5 Lux) per minute. For both PDR and s-PDR assays, we performed four independent experiments (n= 83 wild type and 94 mutant larvae).

### The acoustic evoked response (AR) in light or dark

For the acoustic evoked response (AR), experiments were either started with larvae exposed to light for 90 minutes followed by a dark period of 90 minutes, or to dark first followed by light for the same time periods. Starting at minute 36, larvae were submitted to a recurring one-second sound stimulus of average intensity (= 450 Hz) every 5minutes for a total of 12 stimulations (S1-S12). At stimulation S12 (= minute 90), they were also switched to the alternate light/dark state and then left unstimulated for the following 35 minutes. Starting at minute 126, sounds stimulations were resumed every 5 min. For each assay (light or dark first), we performed three independent experiments (L/D: n= 68 wild type and 62 mutant larvae; D/L: n= 51 wild type and 41 mutant larvae).

### Statistical analyses

We analyzed averaged total traveled distances per larva (with a minimum of triplicate experiments and 24 animals/treatment) in GraphPad Prism (v.8). All results were binned into 1-min intervals and error bars represent the mean standard error of the mean (SEM). Statistical differences between direct comparisons were calculated using multiple t-tests controlling the effect of the correlation among the number of fixed repeated measures. We performed two-way analysis of variance (ANOVA) in graphs when two or more groups were compared simultaneously. Differences with *p* < 0.05 were considered significant (*).

## Acknowledgements

We would like to acknowledge Teresa Nicholson for the gift of antibodies (Ribeye b, Maguk and Cav 1.3). Andrew Seeds and Stefanie Hampel for giving us liberal access to their confocal microscope. Nelson Nancy Cherim, Nelson Santiago Maldonado, Camillo Cangari for preparation of EM sections and Clarissa Del-cueto Carreras for help with EM acquisition.

## Author contributions

MB, LCC and RRM conceived the experiments and wrote the manuscript. LCC, RRM, ASC, JCV and ATT performed all experiments. LCC, RRM, ASC participate in realizing figures of the manuscript. RK and MB performed all EM imaging. GY, BM, SB, OR, and GKV provided intellectual input for technical approaches and maturation of the manuscript. OR, GKV and MB provided intellectual and technical expertise in support of all experiments.

## Materials & Correspondence

Correspondence and material requests should be addressed to

---

**Figure 1. Supplement 1.**
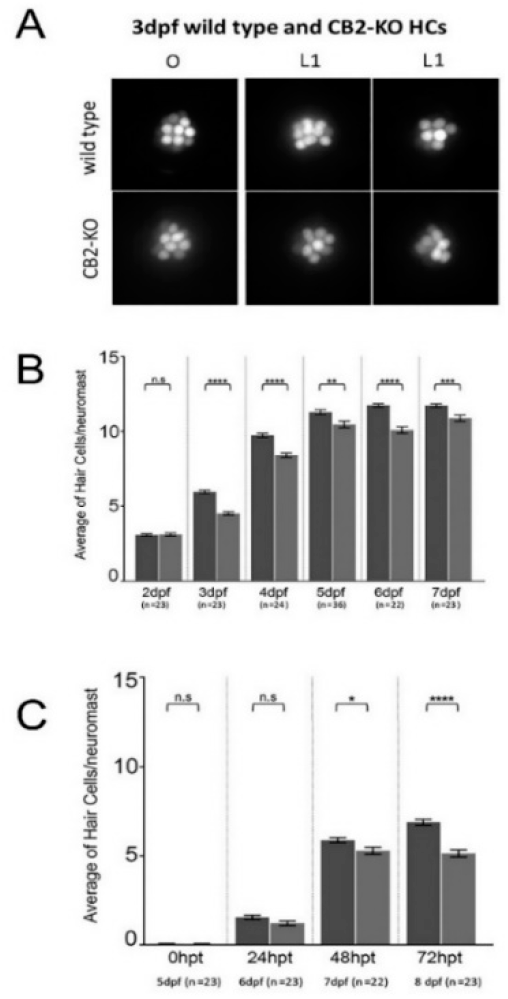
Live imaging of Neuromasts (NMs) after FM 1-43 staining and hair cells (HCs) counts during development and regeneration. (**A**). Magnification of individual cranial NMs (O, left panels) and trunk NMs (L1, middle and right panels) in wild type (top panels) and *cnr2* homozygote mutants (bottom panels). (**B**) Average number of HC/NM at each respective developmental stage (2 to 7dpf) in wild type (dark grey) and *cnr2^upr1/upr1^* (light grey) larvae. (**C**) Average number of regenerated HCs/NM after synchronous ablation of HCs in the LL with a copper treatment (= 0 hour post-treatment, hpt) and subsequent counts performed at 24, 48 and 72hpt in wild type (dark grey) and *cnr2^upr1/upr1^* (light grey) larvae.

**Figure 1. Supplement 2.**
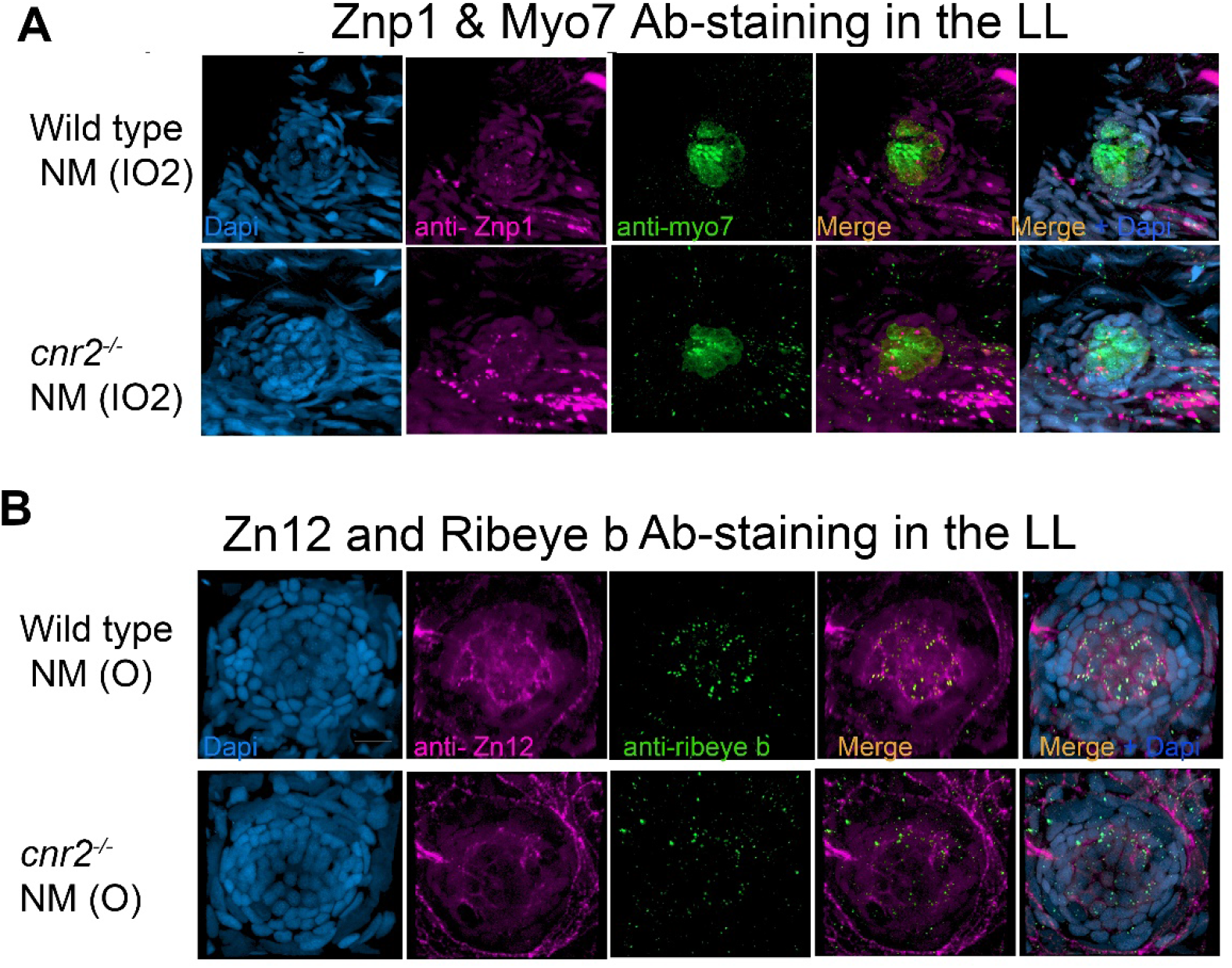
Motor innervation (Znp1) in HCs (Myo7) and sensory innervation (Zn12) with post-synaptic ribbon synapses (Ribeye b) immunolabelling in 5dpf wild type (top panels) and *cnr2^upr1/upr1^* (bottom panels) larvae. (**A**). Top view of wild type (top lanes) and mutant (bottom lane) NMs (IO2) showing motor innervation termination (Znp1, magenta) in HCs stained with myosin 7(Myo7) countered stained with Dapi. (**B**) Top view of wild type (top lane) and mutant (bottom lane) NMs (O) showing sensory innervation (Zn12 in magenta) and presynaptic ribbon synapses in HCs (Ribeye b in green) countered stained with Dapi. Scale bars in A and B = 20 microns. **Supplemental movies:** Figure1-Supplement3-Zn12-Rib_Wt and Figure1-Supplement3-Zn12-Rib-cnr2

**Figure 6. Supplement 1.**
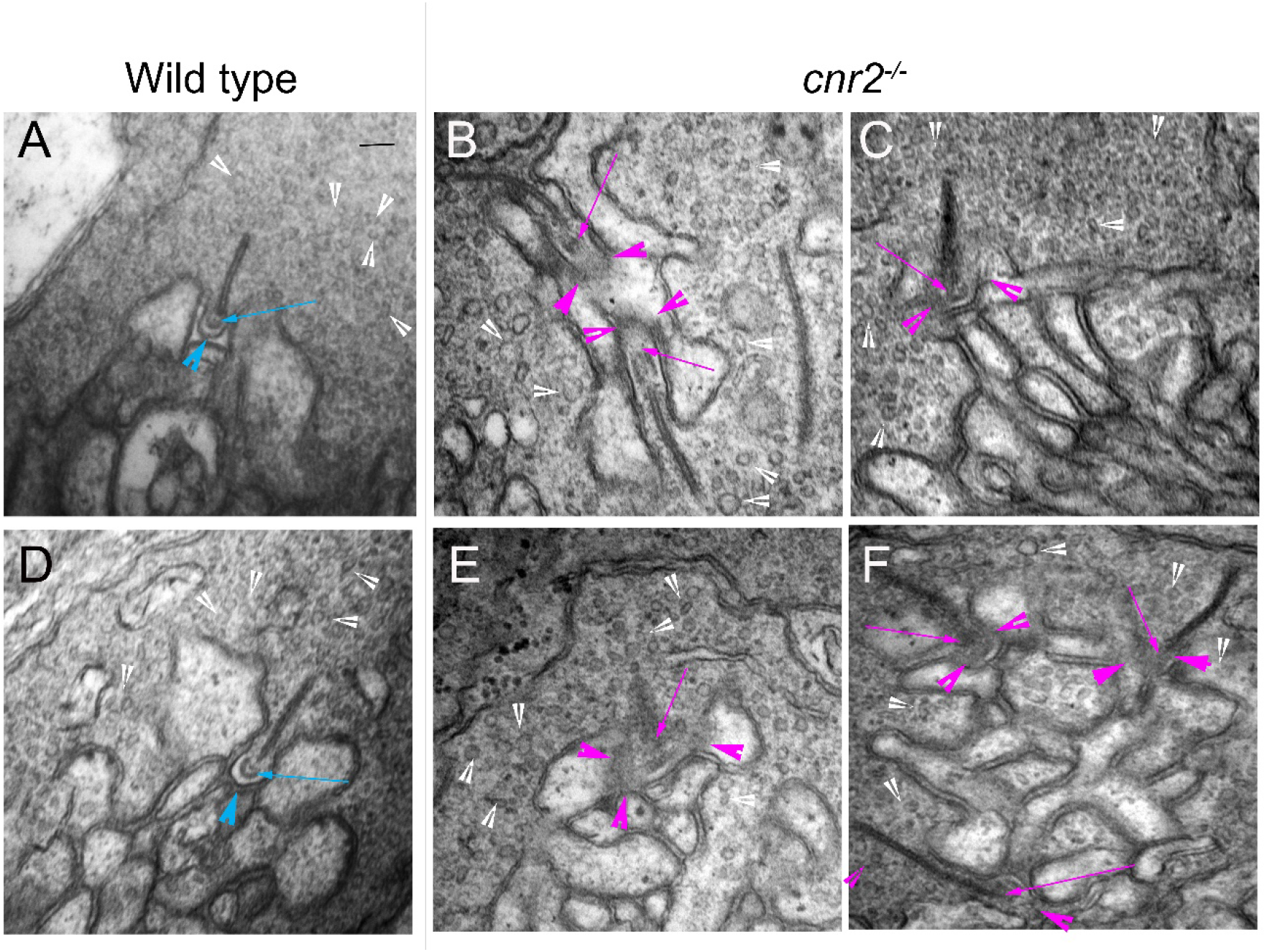
Transmission electron microscopy (TEM) images of cone pedicles showing ribbon synapses in triads in the retina of 5dpf wild type and *cnr2^upr1/upr1^* mutant larvae. (**A** and **D**). In wild type retina, most ribbon synapses have clearly defined arciform densities (blue arrows) in close vicinity to the presynaptic plasma membrane densities (pm, blue arrowheads). They are surrounded by numerous vesicles of ^~^similar size (white arrowheads). (**B**, **C**, **E** and **F**). In *cnr2^upr1/upr1^* retina, most ribbons synapses have poorly defined or incomplete arciform densities (magenta arrows) and presynaptic plasma membrane (magenta arrowheads). The surrounding vesicles appear scarcer and more uneven size (white arrowheads). Scale bar in A representative for all images: = 50 nm.

**Figure 7. supplement 1.**
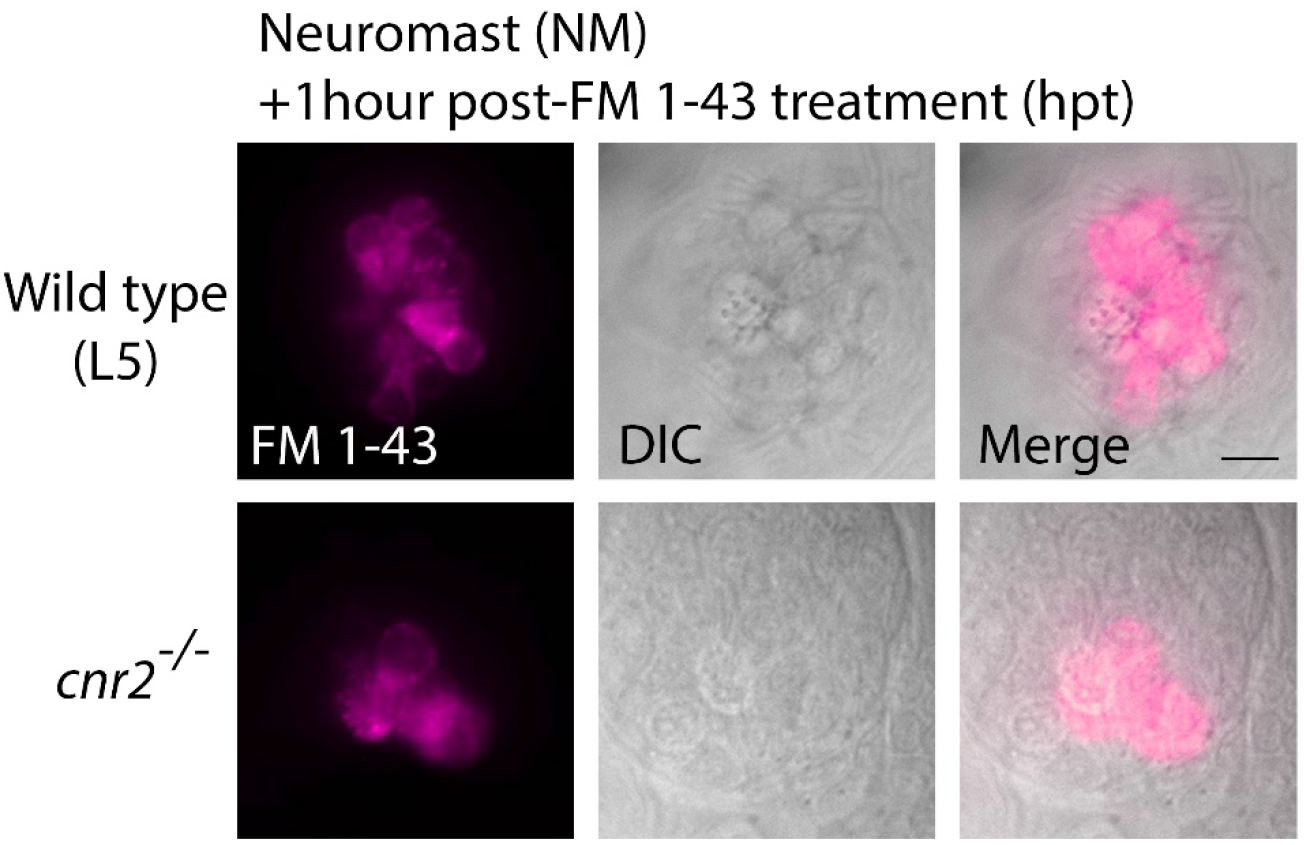
Live imaging of wild type and *cnr2^upr1/upr1^* NMs one-hour post FM 1-43 treatment (hpt). Top views of NM in the pLL (L5) in wild type (top panels) and *cnr2^upr1/upr1^(bottom* panels), showing FM 1-43 (magenta in left and right columns) that penetrated a subset of HCs and dispersed throughout the cell, and brightfield view of the NMs (DIC, center and right columns). Scale bar: = 20 microns.

**Figure 8. Supplement 1.**
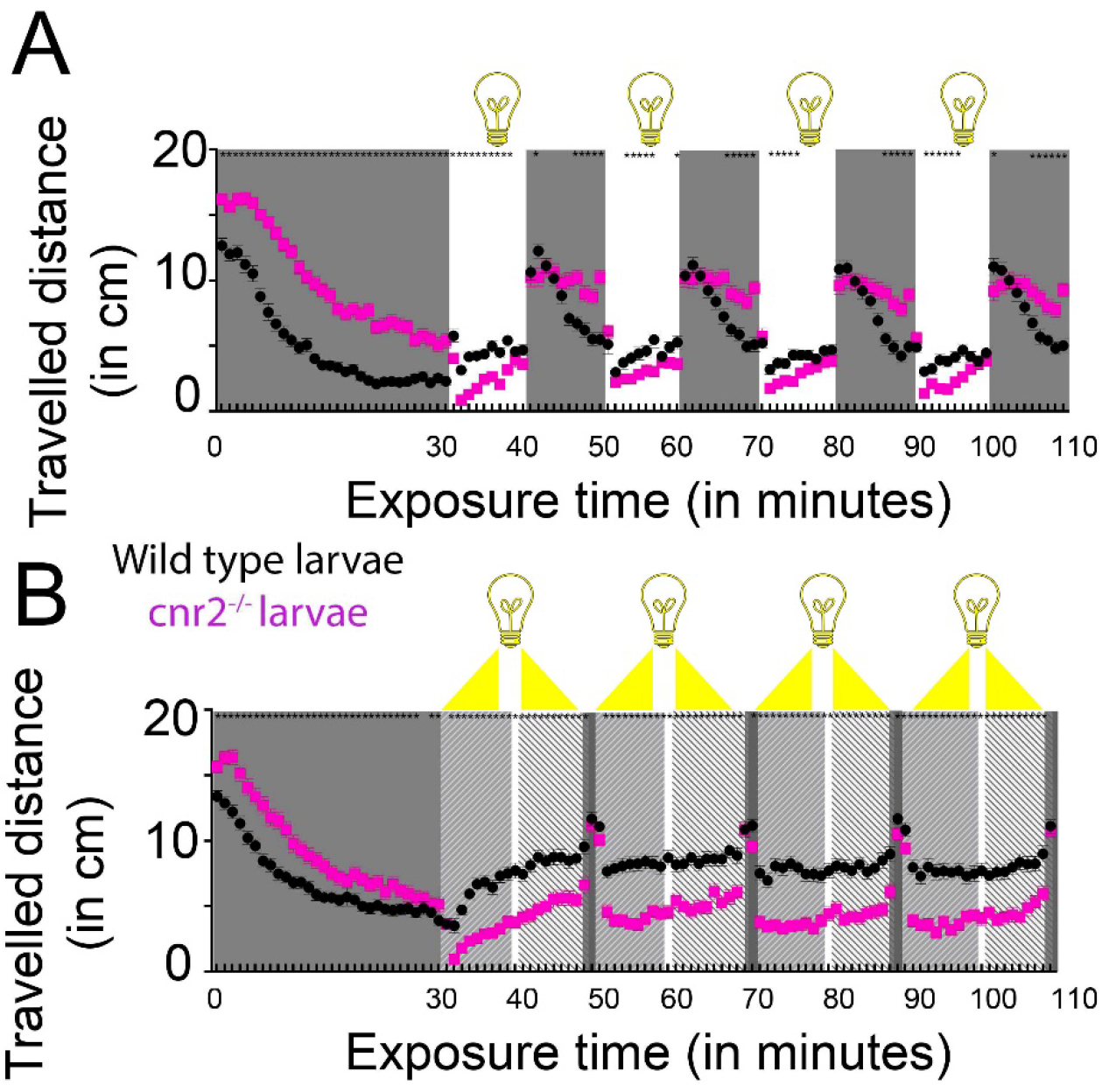
Recording of individual swimming behaviors of 6-day post-fertilization (dpf) wild type and *cnr2^upr1/upr1^* larvae. (**A**) Averaged distance traveled per minute by 6 dpf wild-type (black circles) and *cnr2^upr1/upr1^* (magenta squares) larvae submitted to four successive cycles of 10 min of alternating light periods (white boxes) and dark periods (grey boxes) after a 30-min habituation period to dark. (**B**) Averaged distance traveled per minute by larvae submitted to 4 successive cycles of 10min gradual light intensity increase (0-100%) followed by gradual light intensity decrease (100-0%) after a 30-min habituation period to dark. Error bars represent the standard errors of the mean (SEM) and statistical significance is indicated by* (\**p* < 0.05 and ns is omitted for clarity).

